# Quantification of grapevine yield losses as a function of downy mildew severity on foliage and cluster

**DOI:** 10.1101/2024.02.28.582447

**Authors:** Frédéric Fabre, Lionel Delbac, Charlotte Poeydebat, Marta Zaffaroni

## Abstract

To quantify the relationship between grapevine disease severity and crop losses at the plant scale, we conducted a three-year field experiment at two sites near Bordeaux (France), monitoring mildews and rots on both foliage and clusters. Preliminary analyses revealed that only downy mildew (DM) significantly impacted yield under our experimental conditions. We capitalized on this single-disease dominance to isolate the effect of DM severity (assessed via standardized AUDPC)—on both foliage and clusters—on yield loss at the individual vine scale. At a cluster AUDPC of 5%, yield consistently declined by 24% across all years and sites. Conversely, yield loss at a 5% foliar AUDPC ranged from 9% to 60% depending on the site and year, reflecting a tenfold variation in the damage coefficient. This variability strongly correlated with a moisture risk index computed between the phenological stages of visible inflorescences and pre-ripening berries. Specifically, at 5% foliar AUDPC, the expected yield loss escalated from 9% in a dry season (10th percentile of the index) to 15% in an intermediate season (median), reaching 39% in a wet season (90th percentile). Future research should aim to validate these statistical models across broader climatic, geographic, and agronomic contexts, including the characterization of cultivar-specific responses.

## Introduction

Originating from Eurasia, cultivated grapevine (*Vitis vinifera* subsp. vinifera) has been domesticated and planted for approximately 5 000 years. Several pests and diseases affect grapevine, resulting in the use of high quantities of pesticides. For example, the average treatment frequency index (TFI) for French vineyards in the 2010s was 12.1 with fungicides accounting for 83% to 86% of the total TFI (Fouillet et al., 2022).

Fungicides are mostly applied to control downy mildew (DM) and powdery mildew (PM), the two most important diseases of grapevine (Bois et al., 2017). DM, caused by *Plasmopara viticola*, is one of the most destructive oomycetes in the world (Kamoun et al., 2015). It is considered as the most important disease of grapevine, particularly in European oceanic climate (Bois et al., 2017). DM was first introduced from North America into European vineyards in the 1870s (Millardet, 1881), and subsequently spread to all major grape-producing regions of the world (Fontaine et al., 2021). DM is a polycyclic disease involving during the cropping season primary infections, caused by oospores resulting from sexual reproduction, and secondary infections caused by sporangia resulting from asexual reproduction (Gessler et al., 2011). The relative contribution of primary and secondary infections in the epidemic spread is substantially varying between production situations (Gobbin et al., 2005; Gobbin et al., 2006). While plant infection classically begins on the leaves, DM also infect inflorescences from emergence through to cluster development, until the onset of veraison confers ontogenic resistance (Gindro et al., 2022; Kennelly et al., 2005). This pattern is typical of dual epidemics affecting two plant organs—foliage and inflorescences/clusters—within a single season (Savary et al., 2009). Cluster infections can result from both primary or secondary infections, the latter originating from inoculum produced on the leaves or the clusters. Rain is a major factor fostering DM epidemics as several key steps of primary and secondary infections are strictly dependent on soil wetness, rain splashes or high relative humidity (Bove et al., 2020b; Gessler et al., 2011). The combination of frequent rainfall and moderate temperatures in oceanic climates creates highly favorable conditions for DM epidemics.

PM, caused by the obligate biotrophic fungus *Erysiphe necator*, is also a widespread and destructive disease of grapevines worldwide. As DM, it grows on all green aerial organs and can decrease yield, especially by developing on berries, from flowering to the berry touch stage (Calonnec et al., 2004). Cultivated grapevine can also be frequently infected by two others fungal disease: Botrytis bunch rot (BOT) and black rot (BR). BOT, caused by the necrotrophic fungal pathogen *Botrytis cinerea*, infects mostly inflorescences and ripening berries (Martinez et al., 2005). It could be responsible for important crop losses and qualitative oenological damage (Ky et al., 2012; Steel et al., 2013). BR, caused by the fungi *Phyllosticta ampelicida*, is a foliar and a cluster disease currently becoming increasingly important in Europe following current efforts to reduce fungicide use (Molitor & Beyer, 2014).

The strategies of grapevine protection mainly focus on preventing grape yield losses, while ensuring harvest quality. Accurate estimation of crop losses caused by pathogens, encompassing both yield and quality losses, is central to phytopathological research and, according to some authors, its “raison d’^etre” (James, 1974). In particular, understanding these losses is needed to improve both short-term tactical and long-term strategic management decisions. Pesticide misuse or overuse often result from a poor knowledge of the severity-damage relationship, a key metric used by decision support systems to guide treatments (Savary et al., 2006). Similarly, identifying sustainable strategies for the deployment of resistant cultivars fitting the constraints of real farming systems require mathematical models properly accounting for crop losses (Rimbaud et al., 2021). The initial step toward assessing crop losses involves estimating the attainable yield (*Y_a_*) and the actual yield (*Y*). The actual yield is the yield harvested in a given field, while the attainable yield is the yield that would be harvested without pest attacks over the course of the growing season (Savary et al., 2006; van Ittersum & Rabbinge, 1997; Zadoks & Schein, 1979). The yield losses caused by the considered diseases are then defined as the difference between *Y_a_* and *Y*. Importantly, not all crop losses result in economic losses. A further step is therefore to derive functions that translate crop losses into economic loss (Savary et al., 2018). In viticulture, each commercial context and each production organization (vineyard or winery) has its own definition of the marketable yield (Laurent et al., 2021).

The experimental design of this work was originally set up to assess crop losses due to multiple diseases at individual vine scale. It involves establishing the relationship between injury level and crop loss from field experiments where injury levels are manipulated (Savary et al., 2006). To generate epidemics with contrasting dynamics, fungicides were applied to either suppress or delay the progression of foliar diseases (James, 1974). We monitored the four main grapevine diseases (DM, PM, BOT, and BR) during three successive cropping seasons at two locations in the Bordeaux wine region, under three fungicide management levels. Disease severity was assessed on both foliage and clusters at the individual vine level. Crop losses were quantified in terms of yield (harvested cluster mass) and quality variables linked to grape ripeness (potential alcohol content, pH, acidity, and anthocyanin content). Preliminary statistical analyses showed that, within our experimental design, DM severity was the primary driver of yield at the vine scale, with PM having no detectable additional effect and BOT and BR remaining marginal. Consequently, we model the relationship between DM symptom severity—on both foliage and clusters—and yield losses at the individual vine scale, while retaining PM severity as a control covariate. The analysis of the quality variables is provided as supporting information.

## Materials and methods

### Vineyard sites

The study took place over three growing seasons from 2006 to 2008 at two experimental vineyards located near Bordeaux, Couhins and Latresne, which are approximately 7 kilometers apart (S1 Text). The Bordeaux region is characterized by oceanic climate, typically featuring cool summers and mild winters (for the latitude), with a relatively narrow annual temperature range and few extremes of temperature. Cloudy conditions with frequent precipitations are common, especially in winter. At both locations, the experiments were set up using the *Vitis vinifera* Cabernet-franc cultivar, trained to a double Guyot system. The vine spacing was identical at both sites with values of 1.80 m between rows and 1.10 m within row. Vines at Couhins were planted in 1982, while those at Latresne were planted in 2003. Both sites were equipped with standard weather stations positioned *≤* 300 m from the experimental areas, allowing the daily gathering of minimum and maximum temperature, rainfall and relative humidity.

### Experimental design

At each site, the experimental design consisted of three disease management levels (unprotected, limited disease management, and full disease management) arranged according to a randomized complete block design with six replications or blocks. Each of the 18 individual plots occupied an area of 100 *m*^2^ with 50 plants distributed in five rows. Crop stages were monitored twice a week according to Lancashire et al., 1991. Whereas no fungicide was applied in the unprotected plots, three fungicide treatments were applied against DM, and three against PM, according to cluster development, in the limited disease management plots. In the full disease management plots, 9 fungicide treatments were applied per disease (DM and PM only) between crop stage 16 (sixth leaf unfolded) and crop stage 81 (beginning of ripening) according to a calendar-based schedule. No specific fungicide was used against BR as it was controlled by the side-effects of DM and PM treatments. A single treatment was applied against BOT at stage 85 (softening of berries) in the full disease management level only. Each management level was applied to the same plots during the three cropping seasons studied. The full design and the management practices are described in Savary et al., 2009 and S1 Text, respectively.

### Monitoring of diseases at plant scale

In each plot, disease observations were taken on six vines located at the center of the plot. The percentage of tissue area covered by lesions caused by DM, PM and BR was assessed on the foliage and on the clusters using a visual key that aids positioning the following levels: 0%, 1% and from 5% to 100% with a 5% increment. BOT was assessed on clusters only with the same method. Plants were assessed for the four diseases at seven crop stages: (i) 53 (inflorescences clearly visible), (ii) 55 (inflorescence swelling), (iii) 75 (berries pea-sized), (iv) 77 (begin of berry touch), (v) 79 (all berries touching), (vi) 85 (softening of berries) and (vii) 89 (ripe berries). Crop stages corresponded to averaged crop stage in each site. At each site, all the notations were realized in a single day. On each of the six observed vines per plot, one primary shoot (growing from buds located on the canes) and one secondary shoot (shoots growing from buds located on the vine trunk, at the basis of primary shoots, or on primary shoots below the pruning zone) were chosen at random at the first assessment, and were assessed at the six following observation dates during each growing season. Three leaves were observed on each shoot and at each assessment date: the third, fifth, and last leaf, starting from the apex of the shoot. Accordingly, the observed leaves varied from one observation date to the next. Five clusters were also randomly chosen and assessed on each observed plant. BR was almost absent in all plots (106 zero out of 108 combinations site *×* year *×* management *×* block considered); it was therefore excluded from subsequent analyses.

### Estimation of disease severity at plant scale

The data gathered during disease monitoring were used to estimate disease severity at the individual vine scale. Below, we detail the calculations for mean disease severity and the associated standardized area under the disease progress curve (AUDPC) for DM on the foliage (variable *DA*). The same calculations were performed for each disease (DM, PM and BOT) and organ (foliage, cluster) combination, resulting in the variables *PA*, *DAc*, *PAc* and *BAc*. Let *DSV* (*j, c*) denote the mean disease severity of DM on the foliage for factors combination *j* (*i.e.* an unique combination of site (2 levels), cropping season (3 levels), disease management (3 levels) and block (6 levels)), at crop stage *c* (7 levels). These mean values were estimated in two steps. First, a mean disease severity was calculated for each vine plant at each crop stage by averaging the percentage of tissue area covered with disease lesions of the six leaves observed (three on the primary shoot, three on the secondary shoot). Second, these individual vine severity values were averaged over the six plants observed for each factor combination *j* and each crop stage *c*. The dynamics of DM severity on foliage and clusters across the different site x year x disease management combinations are presented in Figure S1.

Standardized AUDPC summarize disease severity covering different time spans (Madden et al., 2007). Let *t*(*j, c*) denote the day of the year when crop stage *c* is reach for factor combination *j* and *DA*(*j*) the standardized AUDPC of DM severity. *DA*(*j*) was estimated from the mean disease severity values assessed at 7 crop stages *DSV* (*j, c*) and the time interval (in days) between these stages using the trapezoidal method as:

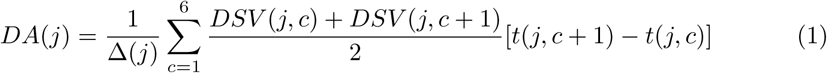

where Δ(*j*) = *t*(*j,* 7) *− t*(*j,* 1) is the duration between crop stages 53 and 89 for factor combination *j*. *DA*(*j*) is expressed in percentage of tissue area. The notations, corresponding variables and model parameters used thereafter are summarized in Table 1.

**Table 1.** Notations used for indices, factors, variables and parameters.

| Type | Notation | Description |
| --- | --- | --- |
| Indice | $j = 1..108$ | Indice over experimental site, year, management and block |
| Indice | $c = 1..7$ | Crop stages 53, 55, 75, 77, 79, 85 and 89 |
| Factor | *Site* | Site (2 levels: Couhins and Latresne, indexed by $s$) |
| Factor | *Year* | Cropping season (3 levels: 2006, 2007 and 2008, indexed by $y$) |
| Factor | *Mng* | Management (3 levels: no, limited and full) |
| Variable | $t(j, c)$ | Day of the year of the crop stage $c$ |
| Variable | $DSV(j, c)$ | Mean severity of DM on leaves (%) |
| Variable | $PSV(j, c)$ | Mean severity of PM on leaves (%) |
| Variable | $DSVc(j, c)$ | Mean severity of DM on clusters (%) |
| Variable | $PSVc(j, c)$ | Mean severity of PM on clusters (%) |
| Variable | $BSVc(j, c)$ | Mean severity of BOT on clusters (%) |
| Variable | $DA(j)$ | Standardized AUDPC of DM severity on leaves |
| Variable | $PA(j)$ | Standardized AUDPC of PM severity on leaves |
| Variable | $DAc(j)$ | Standardized AUDPC of DM severity on clusters |
| Variable | $PAC(j)$ | Standardized AUDPC of PM severity on clusters |
| Variable | $BAC(j)$ | Standardized AUDPC of BOT severity on clusters |
| Variable | $Yield(j)$ | Mean yield (unit: gram) |
| Variable | $MR(s, y, c_f)$ | Moisture risk index between stage 53 and $c_f$ (unitless) |
| Variable | $Z_{lod}$ | Balance resolution used for half-threshold substitution (unit: gram) |
| Parameter | $Y_0(s, y)$ | Mean attainable yield on year $y$ and site $s$ |
| Parameter | $\alpha_{DY}(s, y)$ | DM damage coefficient on yield (foliage) on year $y$ and site $s$ |
| Parameter | $\alpha_{PY}$ | PM control coefficient on yield (foliage) |
| Parameter | $\alpha_{DYc}(s, y)$ | DM damage coefficient on yield (clusters) on year $y$ and site $s$ |
| Parameter | $\alpha_{PYc}$ | PM control coefficient on yield (clusters) |
| Parameter | $\nu_Y$ | Shape of the Gamma distribution of *Yield* |
| Parameter | $a_{DY}$ | Intercept of the post-hoc regression of $\log \alpha_{DY}$ on *MR* |
| Parameter | $b_{DY}$ | Slope of the post-hoc regression of $\log \alpha_{DY}$ on *MR* |

### Assessment of quantitative and qualitative yields

The quantitative actual yield was estimated by the mean yield, *Y ield*(*j*), defined as the average total weight of all clusters harvested from each of the six vines observed for factor combination *j*. A set of qualitative yield variables was assessed on these same six vines using must extracted from 60 berries per plot: potential alcohol content, pH, total acidity, and anthocyanin content (Ribéreau-Gayon et al., 1998). The analysis of these qualitative variables (measurement protocols, statistical analyses, results, and discussion) is presented in S3 Text, allowing us to focus the remainder of the manuscript on the quantitative yield analysis.

### Explanatory statistical analysis

We first investigated the relationship between yield and disease severity at the individual vine scale using generalized linear mixed models (GLMMs) with a Gamma distribution and a logarithmic link. Two models were considered, differing only in the organ on which DM and PM severities were measured. Let *D* and *P*denote the standardized AUDPC of DM and PM on the focal organ—the foliage (*D* = *DA*, *P* = *PA*) in the first model, and the clusters (*D* = *DAc*, *P* = *PAc*) in the second. Letting *BAc* denote the AUDPC of BOT on the clusters, the two models are formulated as:

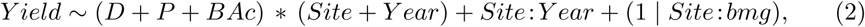

where *∗* denotes main effects and their two-way interactions. The three diseases are thus included both as main effects and as interactions with *Site* and *Y ear*, alongside the *Site × Y ear* interaction. The random intercept over the 3 *×* 6 combinations of disease management and block within sites (hereafter *bmg*) accounts for the fact that the same management levels were applied to the same plots across the three years. The single null yield value was replaced by half the balance resolution (*Z_lod_/*2), the Gamma distribution having no support at zero.

The significance of the fixed effects was assessed with Type II Wald *χ*^2^ tests because several explanatory variables are involved in significant interactions. As a confirmatory analysis, the same models were refitted as linear mixed models on log-transformed yield, log(*Y ield*) (data not shown). Statistical analyses were performed using R version 4.4.1. The GLMMs were fitted with the glmmTMB package (Brooks et al., 2017), the LMMs with the lme4 (Bates et al., 2015) and lmerTest (Kuznetsova et al., 2017) packages, and model residuals were validated using the DHARMa package (Hartig & Lohse, 2020).

### Phenomenological modeling of the relationship between downy mildew severity and yield at the vine scale

As in previous analyses, we assumed a Gamma distribution for *Y ield*(*j*) with a logarithmic link function for its mean:

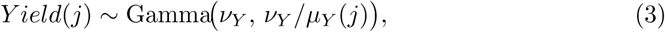

parameterized by a shape parameter *ν_Y_ >* 0 and a rate parameter *ν_Y_ /µ_Y_* (*j*), such that E[*Y ield*(*j*)] = *µ_Y_* (*j*) and the coefficient of variation 1*/√v_Y_* remains constant across the yield range. The single zero-yield record was replaced by *Z_lod_/*2 prior to model fitting, consistent with our earlier approach. Because this half-threshold substitution affected only one of the 108 records, its influence on the parameter estimates was negligible.

### Yield model as a function of DM severity on the foliage

The mean yield combines the attainable yield *Y*_0_(*s, y*) with multiplicative, exponential losses driven by DM and PM severities on the foliage:

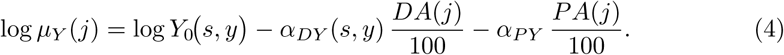

Here, *Y*_0_(*s, y*) is the mean attainable yield for site *s* in cropping season *y*, *α_DY_* is the DM damage coefficient, and *α_P Y_* is a single control coefficient for PM. No interaction between *DA* and *PA* was included: the observed PM gradient was too narrow to identify one (max *PA/*100 *≈* 0.07, cor(*DA, PA*) = 0.23). Thus, *α_P Y_* serves solely to ensure the DM estimate is robust to the presence of PM.

We compared how the DM damage coefficient *α_DY_* varied across sites and cropping seasons. Model *M*_0_ (9 parameters) assumed that DM damage was invariant across sites and seasons by setting *α_DY_* (*s, y*) = *α_DY_ ∀s, y*. In model *M*_1_ (11 parameters), we assumed DM damage depended only on the cropping season (*α_DY_* (*s, y*) = *α_DY_* (*y*) *∀s*). Finally, in model *M*_2_ (14 parameters), DM damage was allowed to vary by both cropping season and site (*α_DY_* (*s, y*)).

### Post-hoc analysis of the moisture dependence of DM damage on the foliage

The comparison of models *M*_0_–*M*_2_ (see Results) indicated that the DM damage coefficient on the foliage varies across cropping seasons and sites. To explore whether this variation is related to moisture conditions, we regressed the site- and year-specific coefficients *α_DY_* (*s, y*) estimated by *M*_2_ on an moisture risk index *MR*(*s, y*), computed between crop stages 53 and 79 for each site *s* and year *y*, using daily rainfall and relative humidity. For each day *d* between crop stages 53 and 79 on a given site *s* and year *y*, let *RF* (*d*) denote the daily rainfall (in *mm*) and *HRm*(*d*) the daily mean relative humidity (in %). First, an ordinal six-class daily moisture risk *MR*(*d, s, y*) was derived from Bove et al. (2020b) and Lalancette (1987) as follows: value 0 if *RF* (*d*) = 0 and *HRm*(*d*) *<* 70; value 1 if *RF* (*d*) = 0 and *HRm*(*d*) *≥* 70; value 2 if 0 *< RF* (*d*) *<* 5 and *HRm*(*d*) *<* 70; value 3 if 0 *< RF* (*d*) *<* 5 and *HRm*(*d*) *≥* 70; value 4 if *RF* (*d*) *≥* 5 and *HRm*(*d*) *<* 70; and value 5 if *RF* (*d*) *≥* 5 and *HRm*(*d*) *≥* 70. Second, *MR*(*s, y*) was computed as the mean of *MR*(*d, s, y*) over all days from crop stage 53 to 79. The annual variation of *MR*(*s, y*) from 1993 to 2022 at the Couhins site is illustrated in Figure S3; its mean and standard deviation are 1.55 and 0.42, respectively.

The regression of *α_DY_* (*s, y*) on *MR*(*s, y*) was carried out on the log scale, consistent with the multiplicative structure of the yield model:

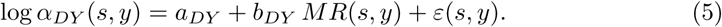

Rather than regressing the six posterior means, we propagated the estimation uncertainty by fitting equation (5) to each posterior draw of *M*_2_: for every MCMC sample, the six values of log *α_DY_* (*s, y*) were regressed on *MR*(*s, y*) by ordinary least squares, yielding a posterior distribution for the intercept *a_DY_*, the slope *b_DY_* and the coefficient of determination. As *MR*(*s, y*) takes only six values clustered in two groups (*≈* 1.08 in 2006 and *≈* 2.08 in 2007 and 2008), this relationship is identified by a two-level contrast rather than by a gradient; it is therefore reported as an exploratory, correlational result rather than as a mechanistic model, and *MR* is not included in the likelihood of the yield model.

### Yield model as a function of DM severity on the clusters

The same structure and the corresponding three specifications for the damage coefficient were fitted to DM severity on the clusters:

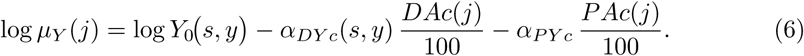

In model *M*_0_ (9 parameters), *α_DY c_*(*s, y*) = *α_DY c_ ∀s, y*; in *M*_1_ (11 parameters), *α_DY c_*(*s, y*) = *α_DY c_*(*y*) *∀s*; and in *M*_2_ (14 parameters), *α_DY c_*(*s, y*) depended on both site and cropping season. Foliage and cluster severities were not entered jointly into the models because *DAc* is largely a downstream consequence of *DA*.

### Derived quantity: relative yield loss

Let *m_s,y_*(*DA*) = *Y*_0_(*s, y*) exp(*− α_DY_* (*s, y*) *DA/*100) denote the PM-free mean yield response to a foliar DM severity *DA ∈* [0, 100]. Here, the attainable yield (*i.e.* yield at zero injury) *Y*_0_(*s, y*) enters the response as a multiplicative scaling factor common to all severities, so that the relative yield did not depend on *Y*_0_(*s, y*). A first-order Taylor expansion for small *DA* then gives the relative loss:

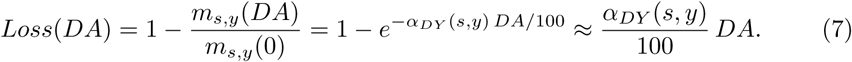

Because *Loss*(*DA*) *∈*[0, 1] is a dimensionless proportion, *α_DY_ /*100 represents the relative yield loss per 1% of standardized AUDPC in the low-severity range; this is the quantity reported throughout the manuscript. The loss thus depends only on the damage coefficient—still possibly varying with site and season through *α_DY_* (*s, y*)—but not on *Y*_0_(*s, y*): it is comparable across sites and seasons despite their different attainable yields, and the choice of a reference *Y*_0_(*s, y*) (e.g., 1000 g) affects only the scale of predicted yields, not the estimated damage. The same definition applies to the clusters, with *m_s,y_*(*DAc*) = *Y*_0_(*s, y*) exp (*− α_DY c_*(*s, y*) *DAc/*100) in place of the foliage response. Reported yields and losses are posterior means of the expected yield E[*µ_Y_* (*·*)], averaged over the joint posterior of (*Y*_0_*, α*).

### Inference of the models with Stan

Model inferences were conducted within a Bayesian framework using Hamiltonian Monte Carlo. The models were implemented in the Stan language via R, interfaced through the cmdstanr package (Stan Development Team, 2023). We used weakly informative priors for all parameters: *Y*_0_(*s, y*) *∼* Normal(1000, 300), *α_DY_* (*s, y*) *∼* Normal(3, 3), *α_P Y_ ∼* Normal(0, 3), *a_DY_ ∼* Normal(1, 1), *b_DY_ ∼* Normal(0, 5), and *ν_Y_ ∼* Gamma(2, 0.1). The same priors were used for the cluster parameters. The distributions for *Y*_0_(*s, y*), *α_DY_* (*s, y*), and their cluster equivalents were strictly lower-bounded at zero, whereas *α_P Y_* was left unbounded so its credible interval could naturally span zero.

We ran 4 chains for 4000 iterations each, including 2000 warmup iterations. Convergence was satisfactory across all models, with *R*^^^ values below 1.01 and no divergent transitions. Model checking and comparison were performed using expected log predictive density (ELPD) via leave-one-out (LOO) cross-validation with Pareto-smoothed importance sampling (PSIS), implemented in the loo package (Vehtari et al., 2017).

## Results

### Multiple disease severity over sites, years and management levels

Substantial variation of climatic conditions was observed between years (Figure S3A). The year 2006, with a mean temperature between crop stages 53 and 79 of 20.9°C in Couhins and 20.3°C in Latresne, was warmer than the years 2007 and 2008 characterized by mean temperatures ranging from 18.4°C (Couhins, 2008) to 18.8°C (Latresne, 2008). During the same period, the years 2007 and 2008 were more humid than 2006. Specifically, the moisture risk index *MR*(*s, y*) was 1.15, 2.21 and 2.16 in Couhins, from 2006 to 2008, respectively (Figure S3B). This variable was systematically lower in Latresne with values 1.02, 1.96 and 2, from 2006 to 2008, respectively.

As expected, the diverse climatic conditions combined with the three disease managements levels led to epidemics of varying magnitude. DM was the predominant disease across all sites and years studied (Figure 1A). The mean (averaged over the block) standardized AUDPC of DM on the foliage ranged from 10% (Latresne, 2006) to 29% (Couhins, 2008) without fungicide control. DM severities were strongly reduced in plots with limited fungicide control, ranging from 0.6% (Latresne, 2006) to 12% (Couhins, 2008), and mostly suppressed under full disease management. The severity of DM on the clusters displayed marked differences between years (Figure 1A, Figure S1). During the cropping seasons 2007 and 2008, the mean standardized AUDPCs of DM on the clusters were very high, with values ranging from 31% (Latresne, 2008) to 66% (Couhins, 2008) without fungicide control and from 17% (Latresne, 2008) to 61% (Couhins, 2008) with limited fungicide control. In contrast, during the cropping season 2006, the mean standardized AUDPC of DM on the clusters remained low, with none exceeding 1.3%. These observations illustrate the complex behavior of dual epidemics on the foliage and the clusters already underlined by Savary et al., 2009.

**Figure 1.**
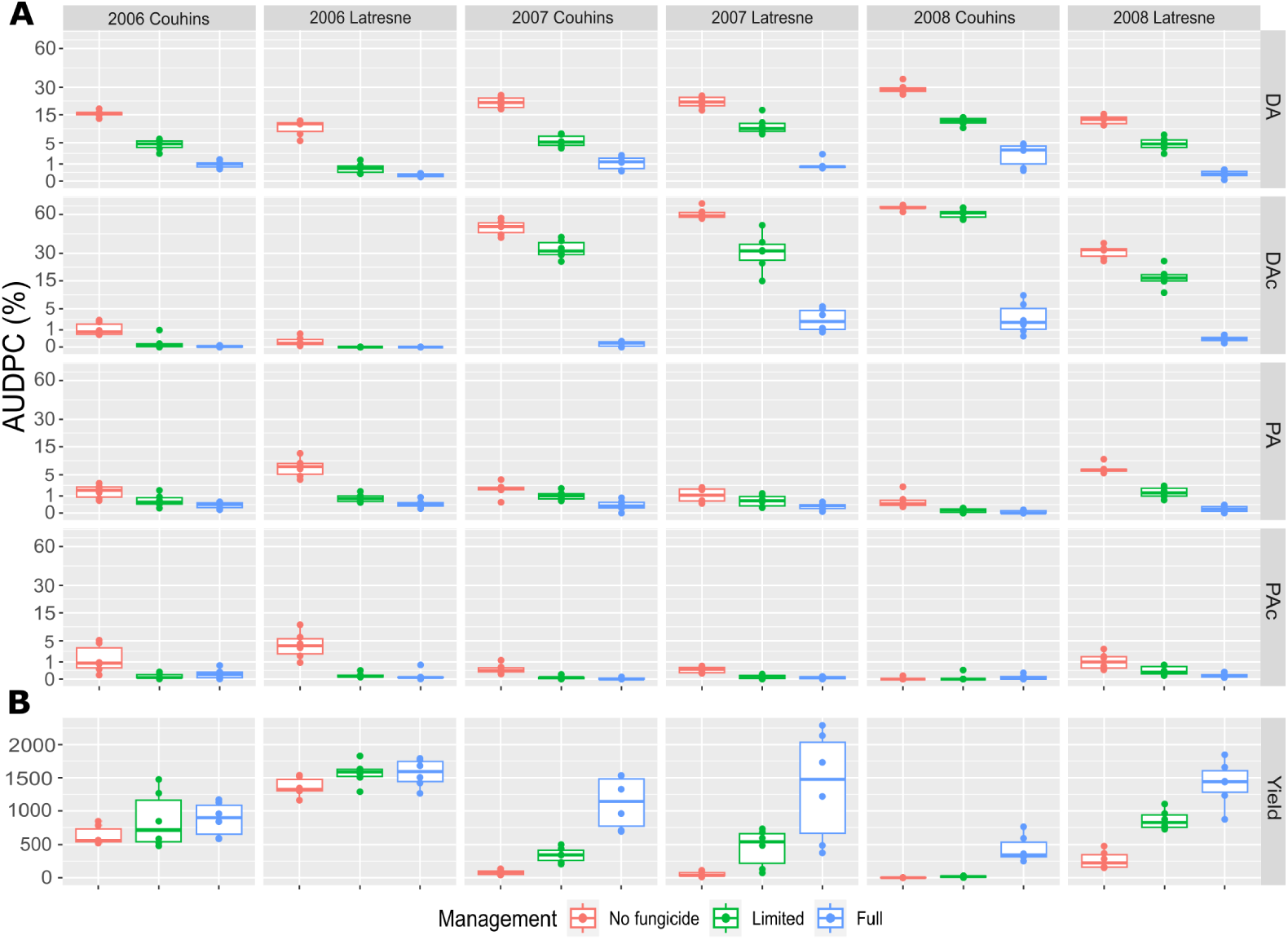
Severity of mildews and yield variables observed at plant scale from 2006 to 2008 in two vineyards under three management levels. (A) The standardized AUDPC summarizes disease severity over time and corresponds to the percentage of tissue area covered by lesions averaged over the whole cropping season. Standardized AUDPC of downy mildew on the foliage (DA, first line) and on the clusters (DAc, second line) and powdery mildew on the foliage (PA, third line) and on the clusters (PAc, fourth line) for each combination of site (Couhins and Latresne), cropping seasons (2006 to 2008) and fungicide managements levels (no, limited and full). Each boxplot is based on the 6 measures (6 replications or blocks) done for a given combination. Note the square root scale of the y-axis. (B) Same as (A) for the total cluster weight per individual vine measured at harvest (in g).

By contrast, BOT epidemics developed very little during the experiments. Even in the plots without fungicide application, the standardized AUDPCs always remained lower than 0.5%. The epidemics of PM displayed an intermediate situation. With fungicide control, the standardized AUDPC was always below 2% both on the foliage and the clusters. Without fungicide, a small PM epidemic was observed on the foliage in Latresne in 2006 (7.3%) and 2008 (6.8%) (Figure 1A, Figure S2). The standardized AUDPC remained below 2% in all other situations. On the clusters, the mean standardized AUDPC of PM severity was always below 2% except at Latresne in 2006 where it reached 4.4 %.

### Relationship between multiple disease severities and yield at the vine scale

The GLMMs with a Gamma distribution described the data satisfactorily, explaining a high proportion of the variance (marginal *R*^2^ = 0.91 and 0.89 for the foliage and cluster models, respectively) and showing almost adequate residual diagnostics (Figure S4).

The corresponding analysis of variance are shown in Table 2. The model incorporating the foliar AUDPCs for DM and PM, alongside the cluster AUDPC for BOT, revealed that yield depended significantly on *Y ear*, *Site*, and their interaction (Figure 1B). It also indicated a strong effect of DM AUDPC, both as a main effect (*χ*^2^ = 327.5) and in interaction with *Y ear* (*χ*^2^ = 54.5). In contrast, the AUDPCs for PM and BOT, as well as their interactions with *Site* or *Y ear*, were not significant, except for a weak main effect of PM (*χ*^2^ = 7.29, *p <* 0.01).

**Table 2.**
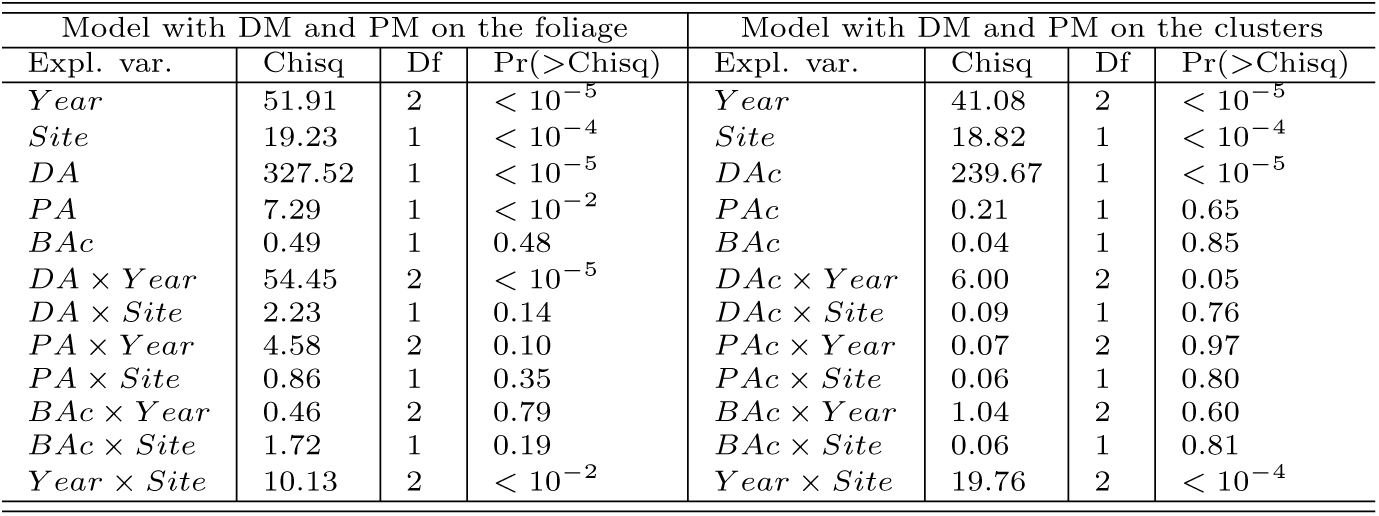
Analysis of variance on the yield (Gamma). Type II Wald *χ*^2^ tests from generalized linear mixed models fitted to *Y ield*, considering as explanatory variables the AUDPC of DM and PM on the foliage (left block) or on the clusters (right block). BOT (*BAc*) was included in both.

When the foliar AUDPCs for DM and PM were replaced by their cluster counterparts, yield again depended on *Y ear*, *Site*, and their interaction, alongside a strong main effect of cluster DM AUDPC (*χ*^2^ = 239.7). A weak *DAc × Y ear* interaction was also detected (*χ*^2^ = 6.0, *p* = 0.05). The corresponding per-year slopes showed a slight difference between 2007 and 2008 (*−*0.052 and *−*0.070 per unit of AUDPC, respectively), while the 2006 slope was estimated over a narrow range of cluster DM severity (AUDPC *<* 1.3%) and was thus poorly identified. This heterogeneity remained small relative to the main DM effect (*χ*^2^ = 6.0 versus 239.7). As shown below, this slight difference did not translate into any improvement in predictive performance, the phenomenological comparison retaining a single, year-invariant damage coefficient.

Finally, refitting both models as linear mixed models on log(*Y ield*) yielded highly similar results, specifically recovering the same set of significant effects (data not shown). However, the Gamma distribution provided a slightly better fit than the log-normal on both foliage (AIC lower by 10.8 points) and cluster (AIC lower by 2.8 points).

### Modeling the relationship between downy mildew severity and yield at plant scale

Previous results indicate that, under our experimental conditions, yield was significantly affected only by DM. While modeling multi-pathogen impacts is feasible, this single-disease dominance provides an ideal context to isolate the effect of DM severity—on both foliage and clusters—on yield loss at the individual vine scale, free from the confounding biases of concurrent epidemics. Consequently, we developed dedicated phenomenological models to specifically characterize this DM–yield relationship.

Let us first consider the three models *M*_0_, *M*_1_ and *M*_2_ fitted to the AUDPC of DM severity on the clusters. No model mis-specification was detected using the PSIS-LOO criteria, all Pareto *k* estimates being good (*k <* 0.5). None of the three models had a significantly better predictive performance than the others, all pairwise differences in ELPD being smaller than their standard error (Table 3). We therefore retained the most parsimonious model, *M*_0_, in which the DM damage coefficient *α_DY c_* does not depend on site or cropping season. This is consistent with the GLMM analysis. Model *M*_0_ fitted the data satisfactorily, with an adjusted *R*^2^ between observed and fitted yield of 0.89 (Fig. 2B). Its estimated parameters are provided in Table 4. The estimated damage coefficient was *α_DY c_* = 5.50 (90% CI [5.05, 5.95]), so that the relative yield loss increases by about 5.5% per 1% of standardized cluster AUDPC in the low-severity range. As expected from the design, the powdery mildew control coefficient was indistinguishable from zero (*α_P Y c_* = 0.10, 90% CI [*−*4.05, 4.02]), confirming that PM had no detectable effect on yield once DM was accounted for.

**Figure 2.**
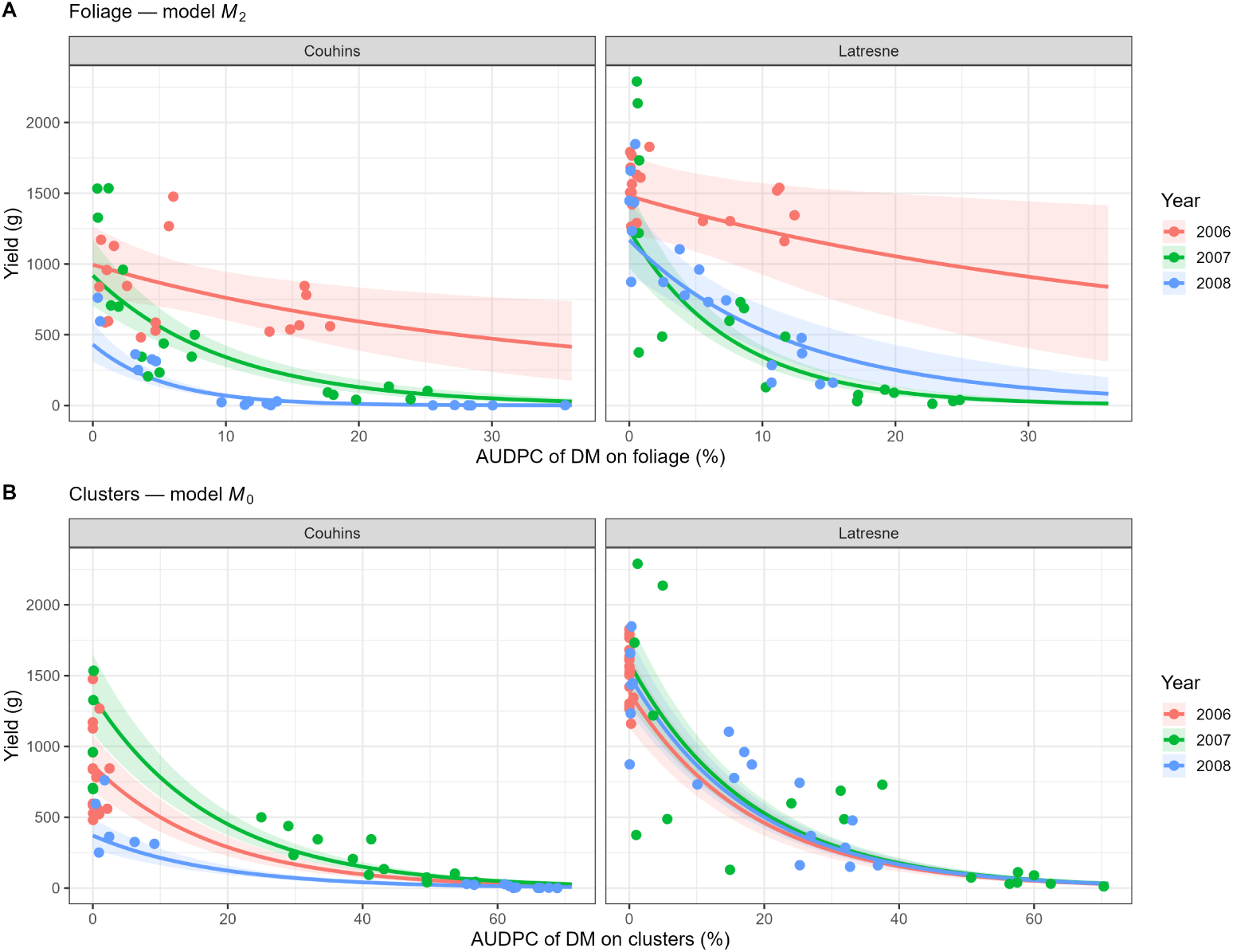
Relationship between downy mildew severity and yield at the individual vine scale, as predicted by the Gamma phenomenological models. (A) Yield as a function of foliar DM AUDPC, fitted by model *M*_2_, where the damage coefficient *α_DY_* (*s, y*) is estimated for each site and cropping season. (B) Yield as a function of cluster DM AUDPC, fitted by model *M*_0_, where the damage coefficient *α_DY c_* is the same across all sites and seasons. Results are displayed separately for the two sites (Couhins and Latresne). Colored lines represent the posterior mean estimated for each cropping season, shaded areas indicate the associated 90% credible intervals, and points represent observed values.

**Table 3.**
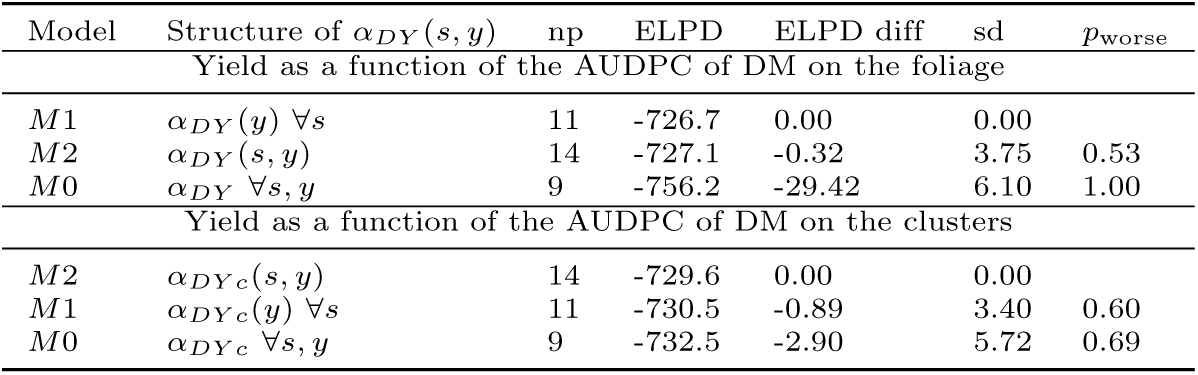
Predictive performance of models relating DM severity to yield at the vine scale. The structural difference between models lies in the specification of the DM damage coefficient, *α_DY_* (*s, y*) (np indicates the number of parameters). Model comparisons are based on the expected log pointwise predictive density (ELPD) and its difference and standard error relative to the best-fitting model (ELPD diff, sd). The *p_worse_* statistic denotes the probability that a given model yields worse predictive performance than the top-ranked model, based on the normal approximation of the ELPD difference; values approaching 0.5 indicate performance indistinguishable from the best model.

**Table 4.**
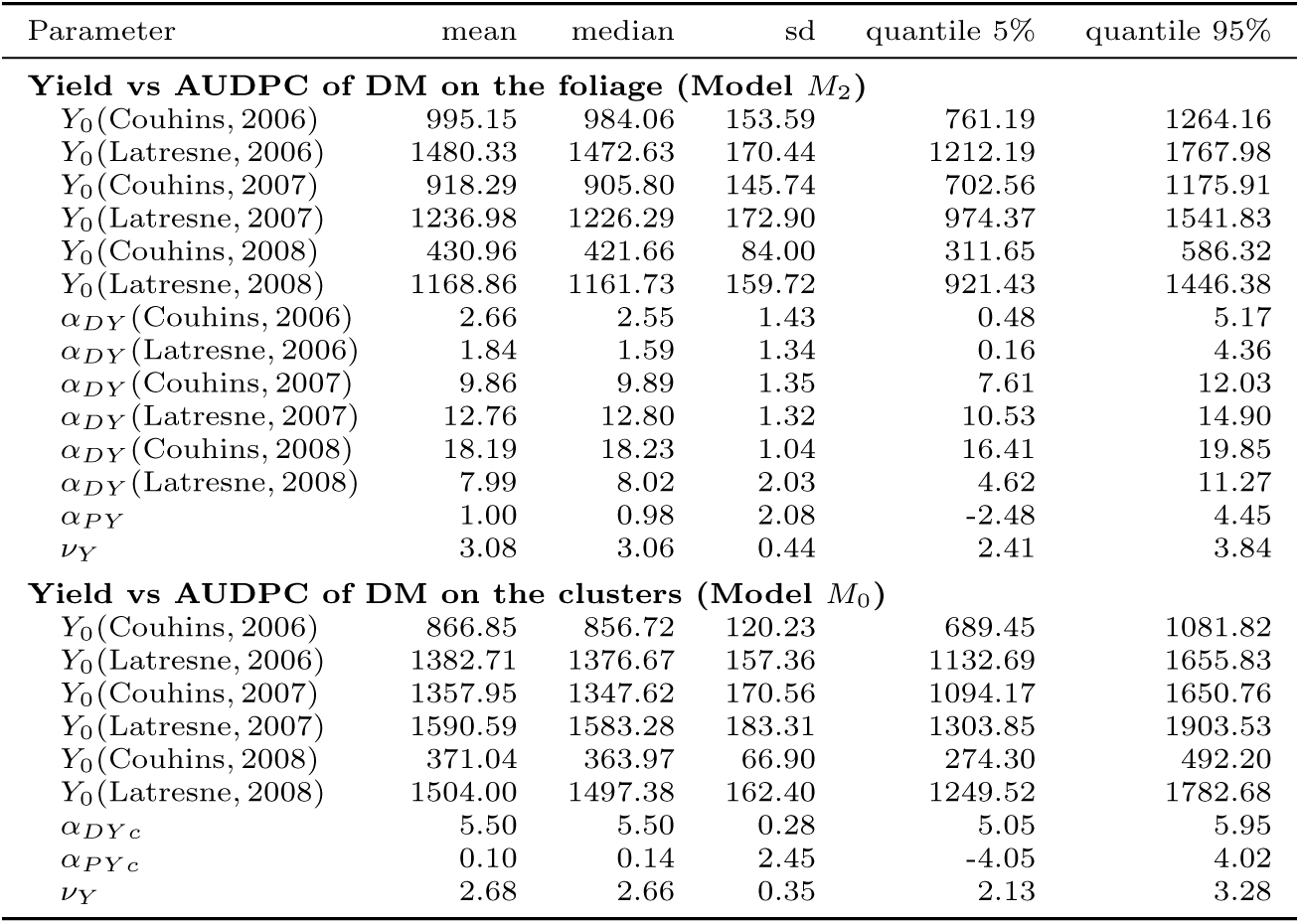
Parameter estimates obtained for the best models fitted to the yield. The first part of the table gives the parameters of model *M*_2_ fitted to the yield as a function of the AUDPC of DM on the foliage, in which the damage coefficient *α_DY_* (*s, y*) depends on both site and cropping season. The second part gives the parameters of model *M*_0_ fitted to the yield as a function of the AUDPC of DM on the clusters, in which the damage coefficient *α_DY c_* is common to all sites and seasons. Model inferences were realized using Hamiltonian Monte Carlo implemented in Stan.

The relationship between the AUDPC of DM on the clusters and the actual yield was predicted for a reference attainable yield of 1000 grams of grapes per vine (Fig. 3D). The mean actual yield declined to 760 g (90% CI [743 g to 777 g]) for a standardized AUDPC of DM on the clusters of 5% (*i.e.* a yield loss of 24%), and to 577 g (90% CI [552 g to 604 g]) for a doubled AUDPC of 10% (*i.e.* a yield loss of 42%). Note that the choice of a reference attainable yield of 1000 grams of grapes per vine corresponds to an interpolation, as this value falls within the range of observed yields across the plots, and facilitates comparisons between sites and years. Predictions can also be made in an extrapolation context, using attainable yield values specific to other production situations.

**Figure 3.**
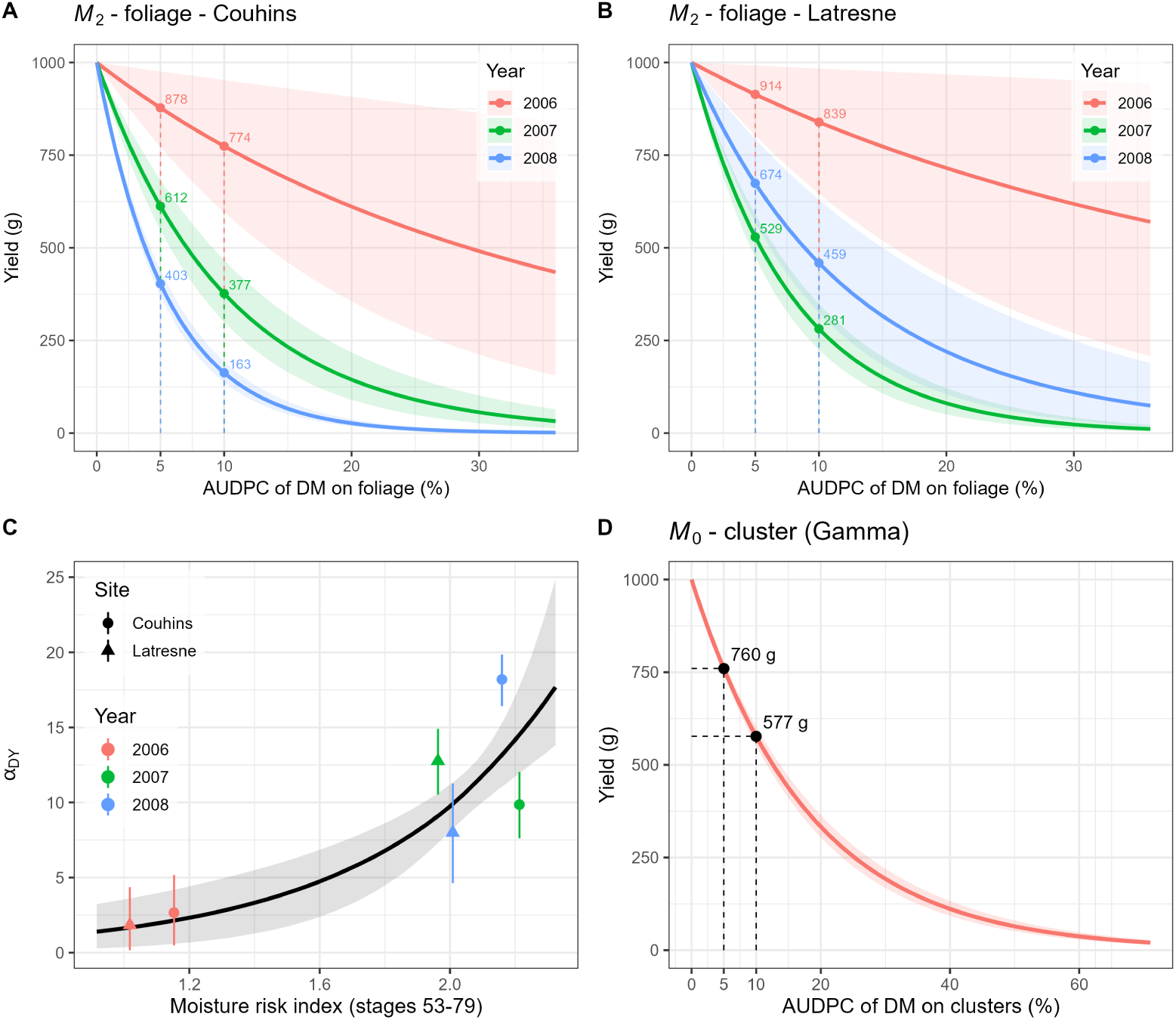
Expected yield losses due to downy mildew, standardized to a reference attainable yield of 1000 g per vine. (A, B) Expected yield as a function of foliar DM AUDPC, estimated by model *M*_2_ for each cropping season, displayed separately for Couhins (A) and Latresne (B). (C) Foliar DM damage coefficient, *α_DY_* (*s, y*), estimated by *M*_2_ (points with 90% credible intervals) versus the moisture risk index (computed between crop stages 53 and 79), overlaid with the post-hoc regression line (posterior median and 90% credible band). (D) Expected yield as a function of cluster DM AUDPC, estimated by model *M*_0_. In panels A, B, and D, colored lines represent the posterior mean expected yields, and shaded areas denote the associated 90% credible intervals; dashed guides and labels indicate the expected mean yield at standardized AUDPC values of 5% and 10%. All yield projections assume a common reference attainable yield of 1000 g to facilitate the comparison of relative losses across sites, seasons, and organs, independent of the actual site- and season-specific attainable yields, *Y*_0_(*s, y*).

We then fitted the three models (*M*_0_, *M*_1_, and *M*_2_) to the AUDPC of DM on the foliage. No model misspecification was detected using the PSIS-LOO diagnostics, as all but one Pareto *k* estimates fell within the reliable range (*k <* 0.7; a single observation reached *k* = 1.04 under *M*_2_, without affecting model selection). Models *M*_1_ and *M*_2_ exhibited indistinguishable predictive performances (ELPD difference smaller than its standard error), and both clearly outperformed *M*_0_ (ELPD difference of *−*29.4, sd 6.1; Table 3). Consistent with the GLMM analysis, this confirms that the foliar DM damage coefficient, *α_DY_* (*s, y*), varies markedly across cropping seasons and sites. We retained *M*_2_, which freely estimates *α_DY_* for each site *×* season combination, as the basis for subsequent post-hoc analyses. This model fitted the data satisfactorily, achieving an adjusted *R*^2^ between observed and fitted yields of 0.91 (Figure 2A).

The parameters estimated for *M*_2_ are given in Table 4. The DM damage coefficient *α_DY_* (*s, y*) ranged from 1.8 (Latresne, 2006) to 18.2 (Couhins, 2008), corresponding to relative yield losses of about 1.8% to 18% per 1% of standardized foliar AUDPC in the low-severity range—a tenfold difference across conditions. The powdery mildew control coefficient was not distinguishable from zero (*α_P Y_* = 1.0, 90% CI [*−*2.5, 4.5]), confirming that PM had no detectable effect on yield once DM was accounted for.

Assuming a 1000 g reference yield, 2006 was consistently the least damaging season. At 5% foliar AUDPC, 2006 showed only modest losses (yielding 878–914 g of grapes across sites), whereas 2008 suffered severe reductions (403–674 g) (Fig. 3A,B). This contrast widened sharply at 10% AUDPC, where the 2008 yield at Couhins decreased to 163 g compared to 774 g in 2006. While 2006 was universally the mildest year, the most destructive epidemic varied slightly by site, occurring in 2008 at Couhins and 2007 at Latresne.

Post-hoc regression of the DM damage coefficients *α_DY_* (*s, y*) of model *M*_2_ against the moisture risk index (calculated over crop stages 53–79) revealed a strong positive association (Figure 3C). The damage coefficient increased from approximately 2 in the driest conditions to nearly 18 in the wettest (slope *b_DY_* = 1.80, 90% CI [1.07, 3.18]; median *R*^2^ = 0.83). Consequently, moisture conditions during cluster susceptibility captured a substantial share of the spatiotemporal variation in foliar DM damage. However, given the limited sample of six site *×* season combinations, this relationship is only an exploratory correlation.

## Discussion

The proper estimation of crop losses caused by diseases is a key knowledge in phytopathology underpinning the design of sustainable crop management systems. The experimental design of this work was originally set up to assess crop losses due to multiple diseases in vineyards under oceanic climatic conditions. Following James (1974) and Savary et al. (2006), it consists in quantifying at the individual vine scale the relationship between injury level and crop losses from field experiments where injury levels are manipulated with fungicide treatments. A first set of statistical analyses revealed that, among the four diseases surveyed, DM was the main driver of yield across the three years and two locations considered. In particular, we did not detect a significant effect of PM severity on yield once DM was accounted for. PM epidemics remained low across most experimental conditions, exceeding 2% severity on grape clusters only at Latresne in 2006 (4.4%). Even in this case, the expected impact of PM on total cluster weight was likely minimal, as berries infected by PM typically experience only a 12% to 20% weight reduction (Calonnec et al., 2004). This epidemiological context, where a single disease predominated, prompted us to model the relationship between DM severity and yield loss at the individual vine scale, as it offered a valuable opportunity to isolate these impacts without the possible confounding due to multi-pathogen interactions.

To our knowledge, this study is the first to precisely quantify the relationship between the severity of DM symptoms on both foliage and clusters and the resulting yield losses at the individual vine level (Figure 2A,B). DM on the clusters reduced yield by 24% at 5% AUDPC and 42% at 10% AUDPC, regardless of year and site (Figure 3D). Foliar DM had a more variable impact, reducing yield at 5% AUDPC by between 9% (Latresne, 2006) and 60% (Couhins, 2008), depending on site and year (Figure 3A,B). Using a dataset collected under northern viticulture conditions by Carisse (2016), who studied DM progress and yield reduction in fields protected against PM and BOT, we fitted a model similar to *M*_0_ (S2 Text). We found a close relationship: at 5% foliar AUDPC, the fitted model predicted a yield loss of 15%, closely matching our own estimate for a season of median moisture (see below) and lending external support to the foliage relationship.

Most previous studies have highlighted the devastating potential of DM by reporting ranges of potential damage, reflecting the collective memory of European viticulture regarding the exceptionally severe epidemics that occurred in Germany, France, and Italy before World War II (Brunet, 1910; Koledenkova et al., 2022). In 2021, Delval et al. (2022) surveyed 21 Walloon winegrowers as vineyards were severely affected by DM under exceptionally wet and cool conditions: eleven growers reported losses above 50% of their optimal yield, and four above 75%. In a simulation study of a Merlot grapevine under 1988–2010 Bordeaux conditions, Leroy et al. (2013) found yield losses above 80% in 14 of the 23 years considered without disease control. Consistently, our exponential models imply that strong epidemics can jeopardize almost the entire harvest: a 50% cluster AUDPC corresponds to a yield loss exceeding 90%, and, under wet-season condition, a 10% foliar AUDPC already exceeds 50% loss. The external dataset of Carisse (2016) supports this trend, with a predicted loss of 28% at 10% foliar AUDPC and of 80% at 50% foliar AUDPC.

Our analysis suggests that yield losses were mainly the consequence of direct losses due to cluster infections by DM. The GLMM and phenomenological analyses consistently indicated that the standardized AUDPC of DM on the clusters explains, with a single coefficient, most of the variability in yield losses; the weak *DAc × Y ear* interaction detected by the GLMM (p-value=0.05) did not improve predictive performance and was therefore not retained. This result aligns with Jermini et al., 2010b, who separated the direct effect of cluster infections from the indirect effect of foliage infections by comparing an “untreated canopy” modality (clusters treated, foliage untreated) with a “standard schedule” (both treated). Direct infections of inflorescences and clusters drastically reduce yield. Before flowering, inflorescences contaminated by *P. viticola* (BBCH 53 to 55) dry up and fall off. Attacks from flowering to early fruit set (BBCH 60 to 71) produce grey rot; both cases result in direct berry losses. Later attacks on still-green berries (BBCH 73 to 79) cause brown rot (Pons et al., 2018). These infected berries contribute almost as much as healthy berries to harvested yield when harvested without sorting, but impair harvest quality as they do not soften (Pons et al., 2018). Consistently, our additional results on qualitative variables (S3 Text) indicate indirect effects of DM on harvest quality after sorting out diseased berries, mainly for potential alcohol content.

Significant differences in the extent of DM damage were observed between cropping seasons. At a 5% foliar AUDPC, model *M*_2_ predicts yield losses ranging from 9% in 2006 (Latresne) to 60% in 2008 (Couhins)—the least and most damaging site–season combinations, respectively (Figure 3A,B). This tenfold variation in the damage coefficient for a given level of foliar severity illustrates the complex interplay between foliage and cluster infections, as previously highlighted by Savary et al. (2009). To disentangle this complexity, we investigated the marked variations in precipitation that characterized these growing seasons. The 2006 season was relatively dry, with a moisture risk index at Couhins of 1.15 (27th percentile of the 30-year historical distribution) and 1.02 at Latresne (7th percentile). In contrast, 2007 and 2008 were substantially wetter, with indices ranging from 1.96 to 2.21 across sites (all *≥* 80^th^ percentile) (Figure S3). A post-hoc regression linked the site- and season-specific damage coefficients, *α_DY_* (*s, y*), to the moisture risk index of Bove and Rossi (2020), computed between phenological stages BBCH 53 (inflorescence clearly visible) and BBCH 79 (all berries touching), the period when inflorescences and clusters are susceptible to DM infection (Gindro et al., 2022; Kennelly et al., 2005). Following the onset of veraison (BBCH 81), neither symptoms nor new infections are usually observed, coinciding with the definitive closure of the stomata that DM uses to penetrate the berries (Gindro et al., 2012). The post-hoc regression revealed a strong positive association (slope *b_DY_* = 1.80, 90% CI [1.07, 3.18]; median *R*^2^ = 0.83). Translating this moisture effect back to our 5% foliar AUDPC baseline, the projected yield loss scales from approximately 9% in a dry season (10th percentile of moisture risk) to 15% in an intermediate season (median), and reaches 39% in a wet season (90th percentile).

This correlation makes no assumption about the origin (primary or secondary) of cluster infections, but implicitly assumes that secondary infections between clusters have little effect on yield. This hypothesis, shared with the process-based model of Bove et al. (2020a), rests on two observations: (i) diseased berries produce sporangia only briefly, and (ii) the transformation of stomata into non-functional lenticels during berry development may inhibit infection and sporulation (Kennelly et al., 2005).

## Conclusions

At the individual vine scale, the relationship between downy mildew severity on clusters and yield loss was described by a single damage coefficient that did not vary across years or sites (24% loss at 5% AUDPC, 42% at 10%). In contrast, the foliar damage coefficient varied about tenfold across site–season combinations. This variation was tentatively associated with growing-season moisture, the site–season combinations falling essentially into two contrasting groups. Deriving robust relationships between foliar DM severity and potential yield loss is nonetheless an important step towards improving grapevine protection: early decisions should ideally rely on assessments of DM on foliage, even though clusters are the main target of protection.

Our yield-loss estimates are assessed at the individual vine level and thus account for possible compensation responses to DM infection at the plant level (Jermini et al., 2010a). However, protection strategies are typically devised at the field level, where compensatory interactions between plants may also occur, particularly if epidemic intensity varies across the field. Another open question is the dependence of the estimated parameters on cultivars: while all *Vitis vinifera* are susceptible to DM, differences in susceptibility between and within cultivars exist (Boso & Kassemeyer, 2008; Gaforio et al., 2015; Paineau et al., 2022). Extending these results will thus require datasets spanning a wider range of agronomic, climatic and geographic conditions to test the proposed models, particularly the moisture–damage relationship currently based on only six conditions (2 sites *×* 3 years). From a modelling perspective, future work could structure the model by crop stages to better account for the decreasing susceptibility of clusters as phenology progresses (Gindro et al., 2022), with stage-specific susceptibility weights estimated directly from the data, and could incorporate inter-annual effects, as the attainable yield in a given year might decrease following DM epidemics in previous seasons (Jermini et al., 2010c).

## Acknowledgments

We thank J.-M. Brustis for his management of the grapevine experiments, the application of pesticide treatments and harvest management. We also thank D. Forget, Head, and all the technical staff of Couhins and Latresne Experimental Stations, INRAE Bordeaux, for allowing these experiments to be conducted and maintain vineyard plots. We also thank Serge Savary and Laetitia Willocquet for setting up the experimental design.

## Funding

This research was funded by INRAE and by the project ANR COMBINE (ANR-22-CE32-0004).

## Conflict of interest disclosure

The authors declare that they comply with the PCI rule of having no financial conflicts of interest in relation to the content of the article.

## Author contributions

Conceptualization: FF, MZ; Data curation: LD; Formal analysis: FF, LD, CP, MZ; Funding acquisition: FF; Investigation: FF, MZ; Methodology: FF, LD, CP, MZ; Writing – original draft: FF, LD, CP, MZ; Writing – review and editing: FF, LD, CP, MZ.

## Data availability

The data as well as R and Stan codes that support the findings of this study are openly available in the dataverse Data INRAE at https://doi.org/10.57745/TVDYAH. Data are currently available via a private URL for review purposes; the dataset will be made publicly available upon publication of the article.

## Supporting information

Supporting information for the article **“Quantification of grapevine yield losses as a function of downy mildew severity on foliage and cluster”** by Frédéric Fabre, Lionel Delbac, Charlotte Poeydebat and Marta Zaffaroni.

### S1 Text

### S2 Text

### S3 Text

The supplementary texts (S1 Text with the experimental design, S2 Text with the analysis of the dataset from the article of Carisse (2016) and S3 Text with the analysis of qualitative yield) are provided in a separate file. The supplementary material also includes one table and six figures, presented below.

---

**Figure S1.**
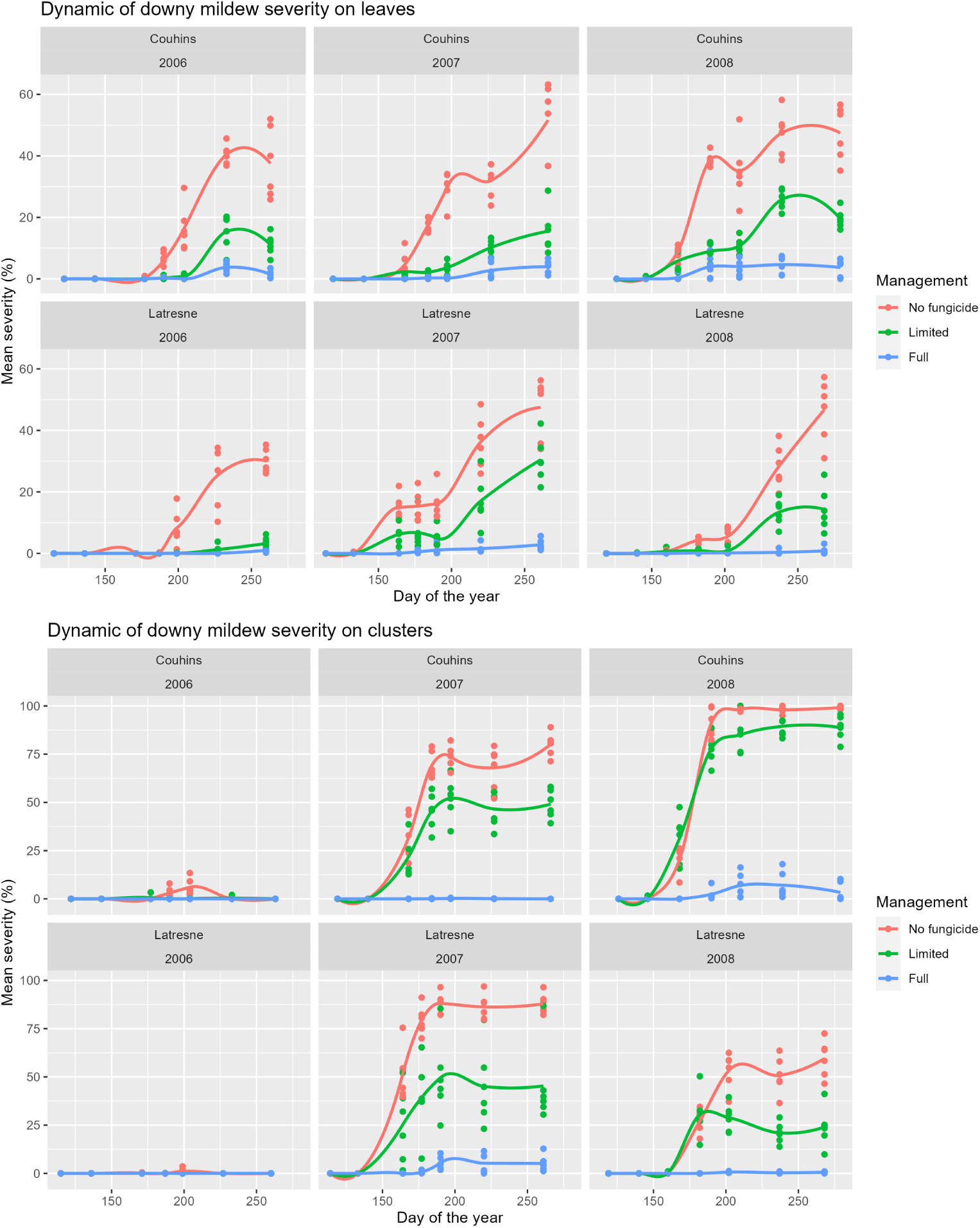
Dynamic of downy mildew severity on leaves and clusters at plant scale. The dynamic of the mean DM severity is plotted as a function of the days of the year corresponding to seven key growth stages on the leaves (top panel) and on the clusters (bottom panel). Points corresponds to the measures realized on the 6 blocks studied for each combination of site (Couhins and Latresne), cropping seasons (2006 to 2008) and fungicide managements levels (no, limited and full). Curves are fitted using a loess method.

**Figure S2.**
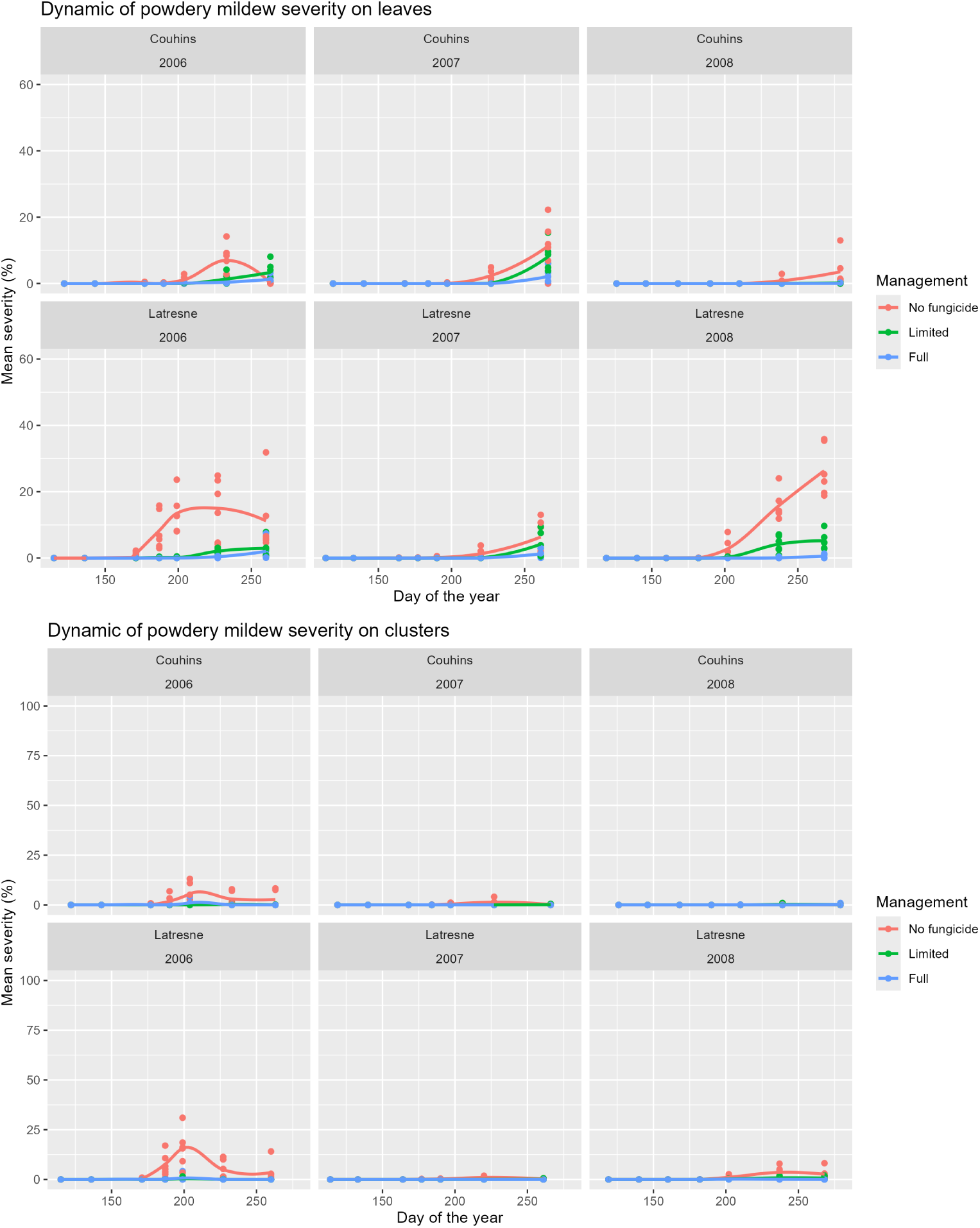
Dynamic of powdery mildew severity on leaves and clusters at plant scale. The dynamic of the mean PM severity is plotted as a function of the days of the year corresponding to seven key growth stages on the leaves (top panel) and on the clusters (bottom panel). Points corresponds to the measures realized on the 6 blocks studied for each combination of site (Couhins and Latresne), cropping seasons (2006 to 2008) and fungicide managements levels (no, limited and full). Curves are fitted using a loess method. The y-axis ranges are matched to those used in the corresponding DM figure to facilitate comparison.

**Figure S3.**
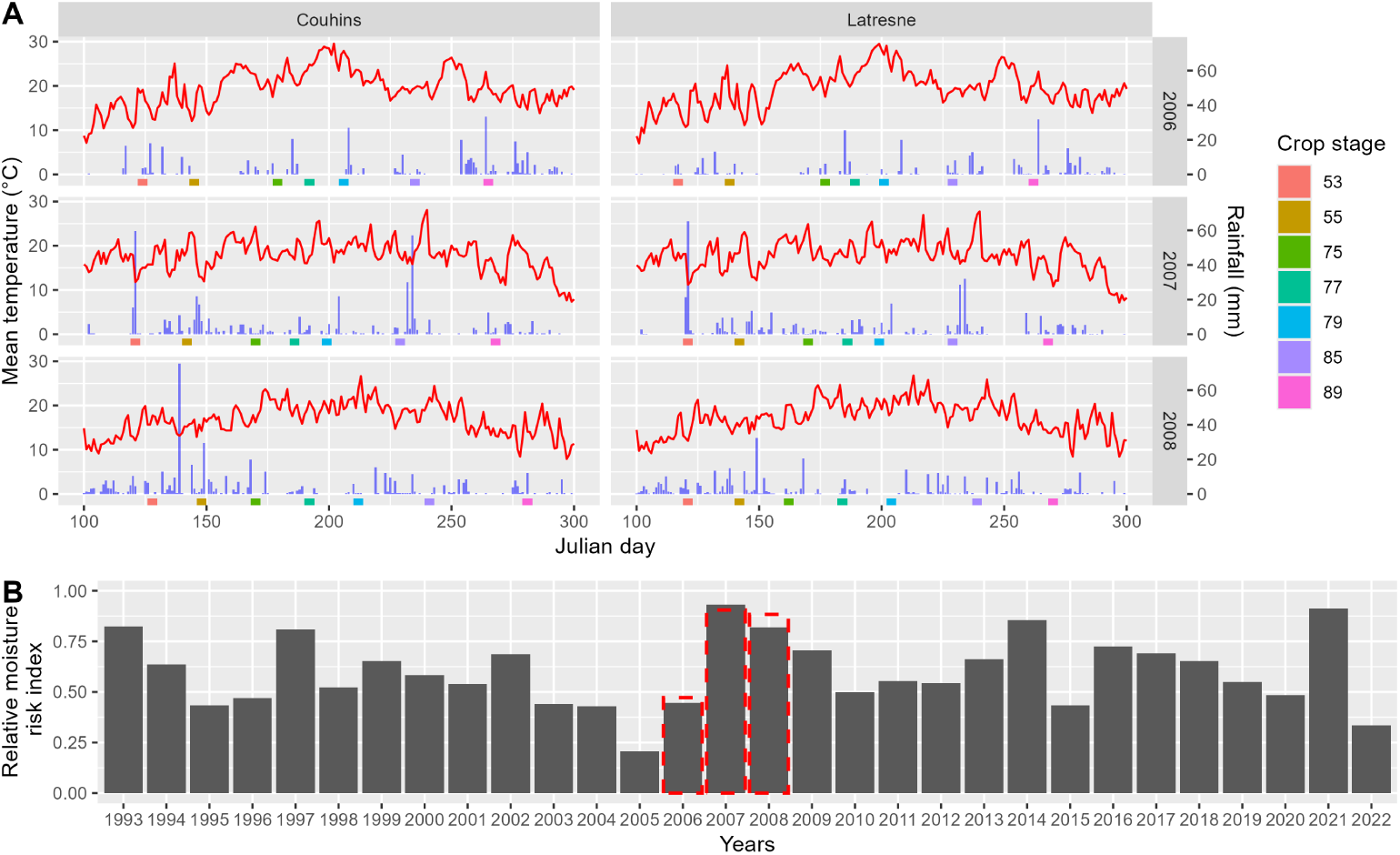
Meteorological data in Couhins and Latresne. (A) Daily mean temperature (red line) and rainfall (blue bar) from day 100 (April 10) to day 300 (October 26), along with the dates of the crop stages where field notations were realized, at the two sites of Couhins and Latresne from 2006 to 2008. (B) The annual value of the moisture risk index is estimated from day of the year 130 (average date of crop stage 53 in our dataset) to day 216 (average date of crop stage 79 in our dataset) in the site of Couhins over 30 years. For the years 2006, 2007, 2008, red dashed bars indicate the value of the moisture risk index for the actual days of arrival of these crop stages.

**Figure S4.**
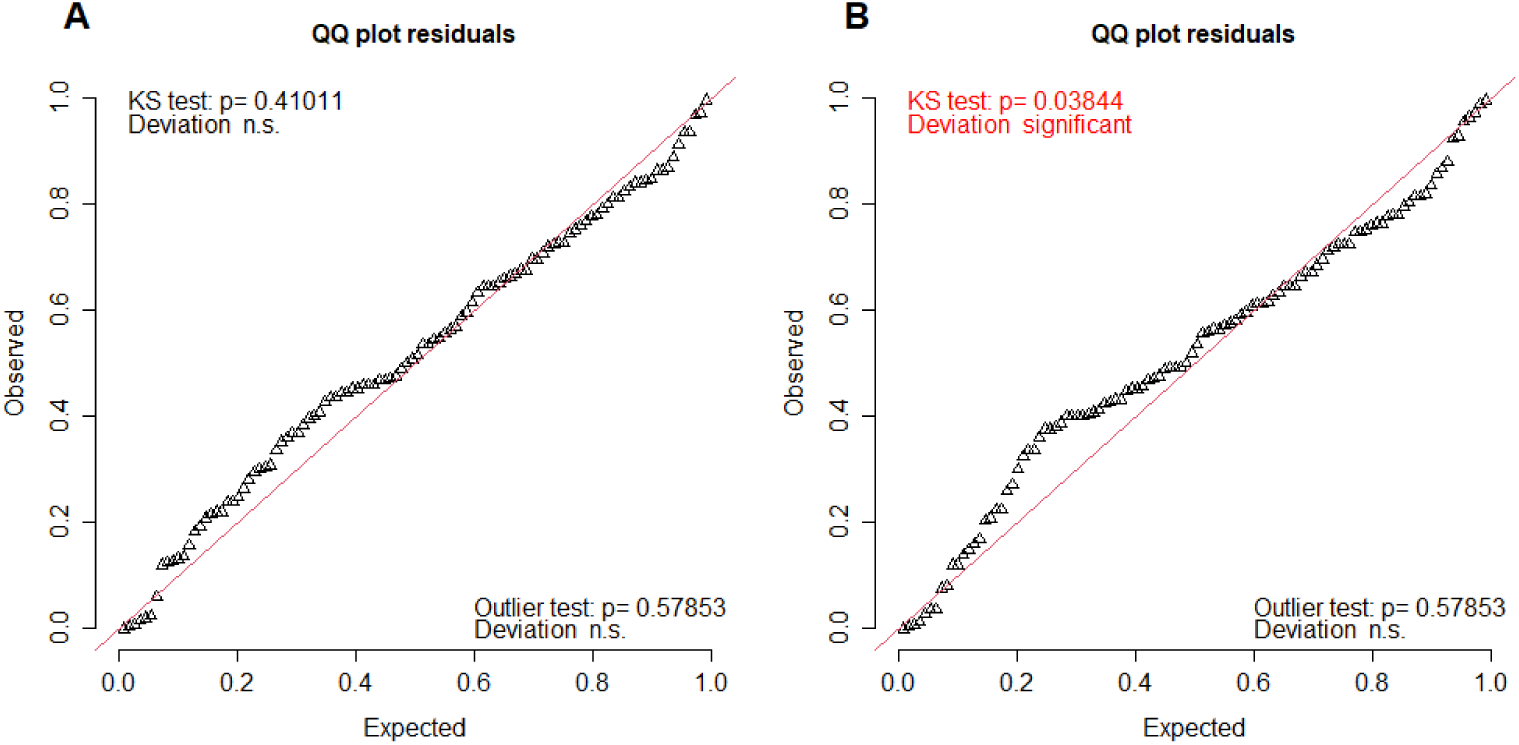
Quantile-quantile plots for the Gamma GLMM fitted to the response variable. *Y ield*. A: Quantile-quantile plot for the GLMM on *Y ield* with the AUDPC of PM and DM on the foliage (variables *PA* and *DA*). B: Quantile-quantile plot for the GLMM on *Y ield* with the AUDPC of PM and DM on the clusters (variables *PAc* and *DAc*). Residuals and plots were computed by simulation using the R package DHARMa.

## Notes

https://doi.org/10.57745/TVDYAH

### Competing Interest Statement

The authors have declared no competing interest.

### Summary of Updates

The word "of" was lacking in the title. Also updated the version of R used (4.4.1 instead of 4.1.3)

