## Supplemental texts for "Quantification of grapevine yield losses as a function of downy mildew severity on foliage and cluster"

S1 Text: Experimental design
S2 Text: Analysis of the dataset from the article of Carisse (2016)
S3 Text: Analysis of qualitative yield

### S1 Text: Experimental design

The details on the experimental design are reported below. The location of the two sites of Couhins and Latresne and the experimental design of the plots are reported in Figure S1.1. The tables report the spraying schedule (Table S1.1), the crop maintenance schedule (Table S1.2) and the characteristics of the compounds used in the spraying schedule (Table S1.3).

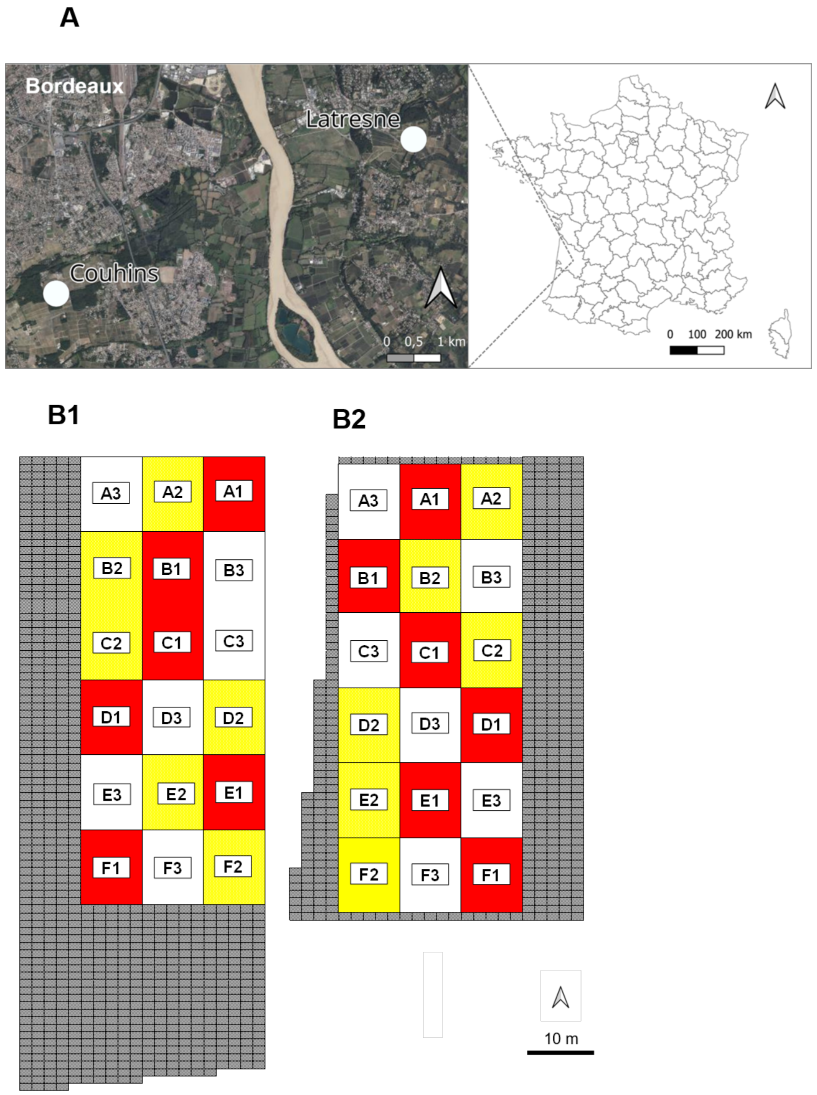


**Figure S1.1. Location of the two sites of Couhins and Latresne and experimental design of the plots.** (A) Location of Couhins and Latresne in the South and East of Bordeaux (France). (B) Experimenatal design of individual plots within the Couhins (B1) and Latresne (B2) vineyards. The colors correspond to the individual plots with no fungicide application (red), limited disease management (yellow) and continuous disease protection (white). In grey, vines not included in the study and managed according to the continuous disease protection. All phytosanitary treatments are applied with a pneumatic knapsack sprayer. Each block is identified by a letter from A to F, with numbers from 1 to 3 corresponding respectively to the preceding colors.

Table S1.1. Spraying schedule in Couhins and Latresne from 2006 to 2008.

| Management | Application no. | Compounds | Dose $ha^{-1}$ | Target | BBCH value's[a] or period | 2006 Couhins Actual BBCH | 2006 Couhins Date | 2006 Latresne Actual BBCH | 2006 Latresne Date | 2007 Couhins Actual BBCH | 2007 Couhins Date | 2007 Latresne Actual BBCH | 2007 Latresne Date | 2008 Couhins Actual BBCH | 2008 Couhins Date | 2008 Latresne Actual BBCH | 2008 Latresne Date |
|---|---|---|---|---|---|---|---|---|---|---|---|---|---|---|---|---|---|
| **Limited disease management** | 1 | tebuconazole, 250 g.liter | 100 g | powdery mildew | 55 | 55 | 5/26 | 55 | 5/19 | 61 | 5/24 | 55 | 5/18 | 55 | 5/28 | 55 | 5/21 |
| | 2 | fosetyl-aluminium, 50% + folpel, 25% | 200 g + 100 g | downy mildew | 61 + 7 days | - | 6/6 | 65 | 6/2 | 69 | 6/7 | 67 | 6/1 | 63 | 6/10 | 62 | 6/4 |
| | | trifloxistrobin, 50% | 62.5 g | powdery mildew | | | | | | | | | | | | | |
| | 3 | fosetyl-aluminium, 50% + folpel, 25% | 200 g + 100 g | downy mildew | Application no.2 + 14 days | - | 6/20 | 72-73 | 6/16 | 73-74 | 6/21 | 74 | 6/15 | 73 | 6/23 | 73 | 6/19 |
| | | tebuconazole, 250 g.liter | 100 g | powdery mildew | | | | | | | | | | | | | |
| | 4 | *Bacillus thuringiensis*, 32,000 UI.mg | 2.4E6 UI | grape moths | Beginning of egg's hatching | 76 | 6/30 | 76 | 6/29 | 76-77 | 7/5 | 75-76 | 6/29 | 76 | 7/7 | 78 | 7/8 |
| | 5 | copper hydroxide, 360 g.liter | 1,530 g | downy mildew | 81 | 81 | 8/10 | - | 8/8 | 81 | 7/31 | 82 | 7/30 | 82 | 8/19 | 83 | 8/18 |
| **Continuous disease protection** | 1 | cymoxanil, 4% + mancozeb, 44.5% | 12 g + 133.5 g | downy mildew | 15-16 | 15 | 5/4 | 15-16 | 4/28 | 17 | 5/3 | 17 | 4/27 | 15 | 5/7 | 15-16 | 5/2 |
| | | wettable sulfur, 825 g.liter | 8,250 g | powdery mildew | | | | | | | | | | | | | |
| | 2 | cymoxanil, 4% + mancozeb, 44.5% | 12 g + 133.5 g | downy mildew | Application no.1 + 10 days | - | 5/15 | 18-19 | 5/9 | - | 5/15 | - | 5/7 | 54 | 5/16 | 54 | 5/13 |
| | | wettable sulfur, 825 g.liter | 8,250 g | powdery mildew | | | | | | | | | | | | | |
| | 3 | fosetyl-aluminium, 50% + folpel, 25% | 200 g + 100 g | downy mildew | Application no.2 + 10 days | 55 | 5/26 | 55 | 5/19 | 61 | 5/24 | 55 | 5/18 | 55 | 5/28 | 55 | 5/21 |
| | | tebuconazole, 250 g.liter | 100 g | powdery mildew | | | | | | | | | | | | | |
| | 4 | fosetyl-aluminium, 50% + folpel, 25% | 200 g + 100 g | downy mildew | Application no.3 + 14 days | 65 | 6/8 | 65 | 6/1 | 69 | 6/7 | 67 | 6/1 | 63 | 6/10 | 62 | 6/4 |
| | | tebuconazole, 250 g.liter | 100 g | powdery mildew | | | | | | | | | | | | | |
| | 5 | fosetyl-aluminium, 50% + folpel, 25% | 200 g + 100 g | downy mildew | Application no.4 + 14 days | - | 6/22 | 72-73 | 6/15 | 73-74 | 6/21 | 74 | 6/15 | 73 | 6/23 | 73 | 6/19 |
| | | quinoxyfen, 250 g.liter | 100 g | powdery mildew | | | | | | | | | | | | | |
| | 6 | fosetyl-aluminium, 50% + folpel, 25% | 200 g + 100 g | downy mildew | Application no.5 + 14 days | - | 7/4 | 76 | 6/27 | 76-77 | 7/5 | 75-76 | 6/29 | 76 | 7/7 | 77 | 7/3 |
| | | quinoxyfen, 250 g.liter | 50 g | powdery mildew | | | | | | | | | | | | | |
| | 7 | lambda-cyhalothrin, 100 g.liter | 17.5 g | grape moths | Beginning of egg's hatching | 76 | 6/30 | 76 | 6/29 | 76-77 | 7/5 | 75-76 | 6/29 | 76 | 7/7 | 78 | 7/8 |
| | 8 | fosetyl-aluminium, 50% + folpel, 25% | 200 g + 100 g | downy mildew | Application no.6 + 14 days | - | 7/18 | - | 7/11 | 79 | 7/19 | 79 | 7/13 | 78 | 7/21 | - | 7/17 |
| | | spiroxamin, 500 g.liter | 300 g | powdery mildew | | | | | | | | | | | | | |
| | 9 | copper hydroxide, 360 g.liter | 1,530 g | downy mildew | Application no.8 + 14 days | - | 8/1 | - | 7/25 | - | 8/2 | - | 7/26 | 79 | 8/4 | 79 | 7/31 |
| | 10 | copper hydroxide, 360 g.liter | 1,530 g | downy mildew | Application no.9 + 14 days | - | 8/11 | - | 8/7 | 82-83 | 8/13 | 83 | 8/6 | 81 | 8/14 | 79 | 8/14 |
| | 11 | pyrimethanil, 400 g.liter | 1,000 g | grey mould | 85 | 86 | 8/30 | 86 | 8/21 | 85-86 | 8/20 | 86 | 8/16 | 85 | 8/25 | 85 | 8/22 |

[a]: Phenological stage of application according Lancashire et al. (1991)

**Table S1.2. Crop maintenance schedule in Couhins and Latresne from 2006 to 2008.**

**Couhins**

| Year | Date | Crop maintenance |
|---|---|---|
| **2006** | March | pruning |
| | 5/5 | tillage |
| | 5/9 | under-row tillage |
| | 5/11 | suckering and shoot thinning |
| | 6/1 | shoot positioning |
| | 6/21 | tip trimming |
| | 6/26 | suckering and shoot thinning |
| | 6/27 | under-row tillage |
| | 7/11 | tip trimming |
| | 7/12 | tillage |
| | 16/8 | tip trimming |
| | 20/8 | under-row tillage |
| | 8/20 | tillage |
| | 9/26 | harvest |
| **2007** | mid February | pruning |
| | 4/23 | tillage |
| | 4/24 | under-row tillage |
| | 5/13 | suckering and shoot thinning |
| | 5/25 | shoot positioning |
| | 6/11 | tip trimming |
| | 6/11 | tillage |
| | 6/15 | under-row tillage |
| | 7/23 | tip trimming |
| | 8/10 | under-row tillage |
| | mid August | tillage |
| | 9/20 | harvest |
| **2008** | February | tillage |
| | end of March | pruning |
| | 5/6 | under-row tillage |
| | 5/26 | suckering and shoot thinning |
| | 6/13 | shoot positioning |
| | 6/13 | tillage |
| | 7/4 | tip trimming |
| | 7/4 | tillage |
| | 7/28 | tip trimming |
| | 8/29 | tip trimming |
| | 8/29 | tillage |
| | mid September | tip trimming |
| | end of September | under-row tillage |
| | 10/7 | harvest |

**Latresne**

| Year | Date | Crop maintenance |
|---|---|---|
| **2006** | February | pruning |
| | end of March | under-row tillage |
| | 4/24 | suckering and shoot thinning |
| | 4/27 | tillage |
| | 6/1 | shoot positioning |
| | 6/15 | tip trimming |
| | 6/26 | suckering and shoot thinning |
| | 6/27 | under-row tillage |
| | 7/18 | tip trimming |
| | 8/10 | tip trimming |
| | 8/10 | tillage |
| | 5/9 | under-row tillage |
| | 9/27 | harvest |
| **2007** | February | pruning |
| | 3/30 | under-row tillage |
| | 4/25 | tillage |
| | 5/15 | suckering and shoot thinning |
| | 6/11 | shoot positioning |
| | 6/11 | under-row tillage |
| | 6/18 | tip trimming |
| | 7/16 | under-row tillage |
| | 7/16 | tillage |
| | 8/20 | tip trimming |
| | mid September | tip trimming |
| | mid of September | under-row tillage |
| | 9/28 | harvest |
| **2008** | February | tillage |
| | March | pruning |
| | 4/1 | under-row tillage |
| | 5/21 | suckering and shoot thinning |
| | 6/2 | shoot positioning |
| | 6/19 | tillage |
| | 6/20 | tip trimming |
| | 7/4 | tip trimming |
| | 7/4 | tillage |
| | 7/18 | under-row tillage |
| | 8/12 | tip trimming |
| | 8/12 | tillage |
| | mid September | tip trimming |
| | end of September | under-row tillage |
| | 10/2 | harvest |

**Table S1.3. Characteristics of the compounds used in the spraying schedule.**

| Compounds | Target [1] | Other Target [1] | Approved for use in France [1, 2] | Persistence (days) [1] | Type of plant action [1] | Mode of Action FRAC / IRAC [3, 4] | Code FRAC / IRAC [3, 4] |
|---|---|---|---|---|---|---|---|
| *Bacillus thuringiensis*, 32,000 UI.mg | grape moths | no | 1984 to still approuved | 12 | contact | midgut | 11A |
| copper hydroxide, 360 g.liter | downy mildew | black rot | 1942 to still approuved | 10 | contact | multi-site | M1 |
| cymoxanil, 4% + mancozeb, 44.5% | downy mildew | black rot, excoriose, rotbrenner | 1978 to 2014 | 10 | penetrating + contact | multi-site: cyanoacetamide-oxime + multi-site: dithiocarbamates and relatives | 27 + M3 |
| fosetyl-aluminium, 50% + folpel, 25% | downy mildew | excoriose | 1979 to 2017 | 14 | systemic + contact | unknown: phosphonate + multi-site: phtalamide | 33 + M4 |
| lambda-cyhalothrin, 100 g.liter | grape moths | beetles, flies,leafhopper, mites | 1988 to 2015 | 14 | contact | nerve and muscle | 3A |
| pyrimethanil, 400 g.liter | grey mould | OTA-producing fungus | 1994 to still approuved | 14 | contact | D1: AP fungicides | 9 |
| quinoxyfen, 250 g.liter | powdery mildew | no | 2000 to 2014 | 14 | systemic | E1: Aza-naphthalenes | 13 |
| spiroxamin, 500 g.liter | powdery mildew | no | 1998 to still approuved | 14 | systemic | G2: Amines (morpholines) SBI Class II | 5 |
| tebuconazole, 250 g.liter | powdery mildew | black rot, rotbrenner | 1994 to still approuved | 14 | systemic | G1: DMI (SBI class 1) | 3 |
| trifloxystrobin, 50% | powdery mildew | black rot, excoriose, rotbrenner | 2002 to 2014 | 14 | penetrating | C3: QoI | 11 |
| wettable sulfur, 825 g.liter | powdery mildew | anthracnose | 1961 to still approuved | 10 | contact | multi-site | M2 |

1) Index phytosanitaires Acta, Acta-publications, Paris, France, 1961 to 2024

2) Spinosi, J., Chaperon, L., Jezewski-Serra, D., & El Yamani, M. (2017). Matphyto, une approche de l'exposition aux pesticides par l'utilisation des matrices cultures expositions : Cas des pesticides arsenicaux. Archives des Maladies Professionnelles et de l'Environnement, 78(5), 472. https://doi.org/10.1016/j.admp.2017.07.010

3) FRAC List of fungicide common names - 2016 (https://www.frac.info/knowledge-database/downloads)

4) IRAC Mode of action classification scheme (https://irac-online.org/?s=)

### S2 Text: Analysis of the dataset from the article of Carisse (2016)

Carisse (2016) monitored DM from 2009 to 2011 in vineyard plots planted with the grape cultivars Chancellor, Vidal and Seyval Blanc. They measured the standardized AUDPC of DM and yield, expressed as the total weight of marketable clusters per vine. The data provided in this article (more specifically in tables 1, 2 and 3; $n = 36$) were used to fit a single-disease version of model $M_0$, with a single attainable yield $Y_0$ (depending on neither cultivar nor cropping season). As this dataset was collected in plots protected against all diseases except DM, no PM control covariate was included.

The estimated parameters are given in Table S2.1. The model fitted the data satisfactorily (Figure S2.1A), with an adjusted $R^2$ of 0.73 between observed and fitted yields on the log scale on which the model is defined. The estimated attainable yield was $Y_0 = 1953$ g (90% CI [1817, 2093]) and the DM damage coefficient $\alpha_{DY} = 3.23$ (90% CI [2.60, 3.83]). Standardized to a 1000 g reference attainable yield, this corresponds to a predicted yield loss of 15% at 5% foliar AUDPC, 28% at 10%, and 80% at 50% (Figure S2.1B). The loss at 5% AUDPC closely matches our own estimate for a season of median moisture (15%), providing external support to the foliar DM–yield relationship derived from the main experiment.

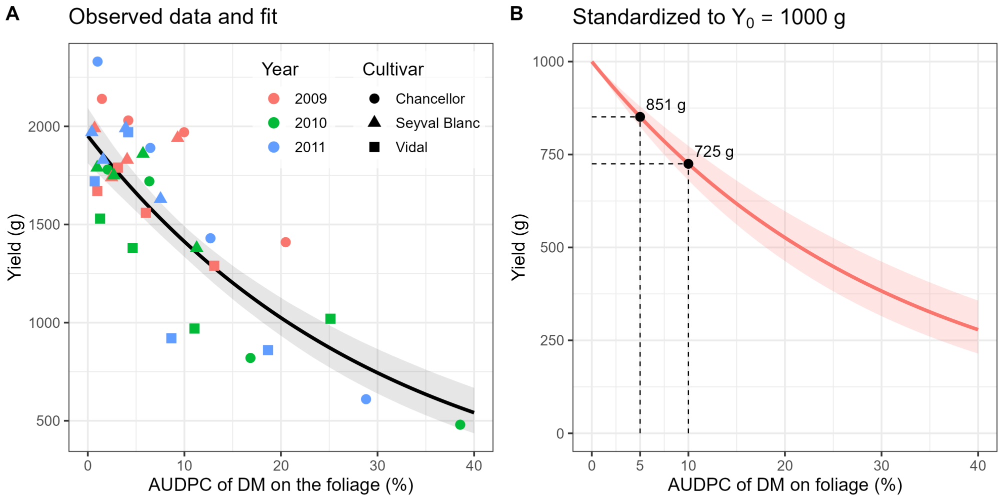


**Figure S2.1. Downy mildew severity–yield relationship fitted to the dataset of Carisse (2016) collected under northern viticulture conditions.** A Gamma model analogous to $M_0$ (but with DM only, no PM covariate) was fitted to fields protected against powdery mildew and *Botrytis*. (A) Observed yields (points, by year and cultivar) and fitted relationship at the estimated attainable yield ($Y_0 = 1953$ g); the line is the posterior mean and the shaded area the 90% credible interval. (B) The same relationship standardized to a reference attainable yield of 1000 g. Dashed guides mark the predicted mean yield at foliar AUDPC values of 5% and 10%.

**Table S2.1.** Parameter estimates for the model fitted to the Carisse, 2016 dataset (Gamma observation model). $Y_0$ is the single attainable yield (g) and $\alpha_{DY}$ the DM damage coefficient on the foliage. Predicted yields are given at a 1000 g reference for a foliar AUDPC of 1, 5, 10 and 50%.

| Parameter | mean | median | sd | quantile 5% | quantile 95% |
|---|---|---|---|---|---|
| $Y_0$ | 1951.199 | 1950.857 | 85.308 | 1812.142 | 2092.877 |
| $\alpha_{DY}$ | 3.224 | 3.232 | 0.388 | 2.575 | 3.846 |
| $\nu_Y$ | 27.036 | 26.582 | 6.134 | 17.655 | 37.776 |
| Yield at 1% DM | 968.281 | 968.199 | 3.756 | 962.266 | 974.576 |
| Yield at 5% DM | 851.279 | 850.792 | 16.533 | 825.043 | 879.180 |
| Yield at 10% DM | 724.950 | 723.846 | 28.207 | 680.695 | 772.958 |
| Yield at 50% DM | 203.307 | 198.716 | 40.486 | 146.138 | 275.918 |

### S3 Text: Analysis of qualitative yield

#### Materials and Methods

**Assessment of qualitative yield**. A set of qualitative yield variables were assessed on the same six plants observed for quantitative yield for each factor combination $j$. Before harvest, 10 healthy berries were sampled from 6 clusters per plot (one cluster per surveyed vine) for a total of 60 berries per plot. The 60-berries sample was crushed and the must analyzed for characterizing grape quality. Since only berries without visible symptoms were used in this sampling, the grape quality measurements obtained represent the quality expected when diseased berries are removed during harvesting. Measuring grape quality, a complex task typically dependent on the type of wine produced, involves assessing the must through a combination of various chemical analyses (Pérez-Álvarez et al., 2021; Ribéreau-Gayon et al., 1998; VanderWeide et al., 2021).

Four oenological indicators were measured on the must (Ribéreau-Gayon et al., 1998). First, the total soluble solids contained in the must were determined as °Brix using a hand refractometer with temperature compensation (Ribéreau-Gayon et al., 1998). Then °Brix was transformed in potential alcohol content $AlDg(j)$ (unit: % vol) using a conversion table (Figure S3.1A). Secondly, the pH $pH(j)$ (unitless) was measured using a pH electrode (Figure S3.1B). Thirdly, total acidity $Acid(j)$ (unit: g $H_2SO_4$/l) was assessed as the equivalent of grams of sulphuric acid per litre (unit: g $H_2SO_4$/l). This value was measured by titration with NaOH 0.1N and with indicator (bromothymol blue). Finally, the anthocyanin content $Antho(j)$ (unit: mg/l) was measured using the Glories method. The absorbance of solution of pH = 1 (0.1 N HCl) was then measured at 520 nm with an UV/VIS spectrophotometer (Saint Cricq de Gaulejac et al., 1998). These quality variables were not available in 7 plots (hence $n = 101$ for these four variables).

**Statistical analysis**. We investigated the relationship between qualitative yield (response variables $AlDg$ and $pH$) and the AUDPC of the four diseases at plant scale by fitting linear mixed models (LMM). The same model structure as the one considered for the quantitative yield $Yield$ was used, replacing the Gamma GLMM by a Gaussian LMM as $AlDg$ and $pH$ are continuous, approximately symmetric responses. We omitted the other two quality variables ($Antho$ and $Acid$) since they were strongly correlated with $AlDg$ ($r = 0.52$) and $pH$ ($r = -0.61$), respectively (Figure S3.2).

Analyses were performed with R version 4.1.3. The LMMs were fitted with the lmerTest package (Kuznetsova et al., 2017); residuals were checked with the DHARMa package (Hartig & Lohse, 2020).

#### Results and discussion

The LMMs used to analyse the potential alcohol $AlDg$ fitted the data in a satisfactory manner (diagnostic tests of residuals, Figure S3.3 A-D, adjusted $R^2 = 0.8$ when considering the AUDPC of DM and PM on the foliage). As for yield, $AlDg$ depended on a strong interaction between $Year$ and $Site$ (Table S3.1) and on highly significant effects of the AUDPC of DM on the foliage alone and in interaction with $Year$. The potential alcohol content decreased with increasing AUDPC of DM, showing a steeper decline in 2006 compared to 2007 and 2008 (Figure S3.4 A). The effect of PM and BOT were not significant alone but slightly significant effect of interactions between $PA \times Site$ (p-value=0.05) and $BAc \times Year$ (p-value=0.02) existed (Table S3.1). Unlike the analysis on $Yield$, replacing the AUDPC of PM and DM on the foliage by their counterparts on the clusters did not simplify the picture. DM, alone and in interactions with $Year$ or $Site$, was associated to highly significant effects on $AlDg$. A significant effect of PM alone was also detected (p-value=0.03) while BOT was no more associated to any significant effect.

The LMMs used to analyse the response variable $pH$ also fitted the data in a satisfactory manner (diagnostic tests of residuals, Figure S3.5 A-D) but with a lower adjusted $R^2$ of 0.58 when considering the AUDPC of DM and PM on the foliage. Highly significant effects of the $Year$ and $Site$, along with their interaction, were evidenced on the pH. The pH increased slightly but significantly with the AUDPC of DM on the foliage (Figure S3.4 B; Table S3.1, p-value=0.03). The pH increased by only 0.1 from low to high DM severity values. When replacing the AUDPC of PM and DM on the foliage by their counterparts on the clusters, only $Year$, $Site$ and their interaction significantly affected the $pH$. The AUDPC of DM on the clusters did not exhibited a significant effect on the $pH$.

**Table S3.1. Analysis of variance on the potential alcohol content and the pH.** The LMMs fitted considered as explanatory variables the AUDPC of DM and PM on foliage (top block) or clusters (bottom block).

| Resp. var. | $AlDg$ | | | $pH$ | | |
|---|---|---|---|---|---|---|
| LMMs fitted with the AUDPC of DM and PM on the foliage | | | | | | |
| Expl. var. | NumDf | F value | Pr(>F) | NumDf | F value | Pr(>F) |
| $Year$ | 2 | 2.86 | 0.06 | 2 | 16.57 | $< 10^{-5}$ |
| $Site$ | 1 | 9.24 | $< 10^{-5}$ | 1 | 14.17 | $< 10^{-3}$ |
| $DA$ | 1 | 68.12 | $< 10^{-5}$ | 1 | 4.72 | 0.03 |
| $PA$ | 1 | 0.96 | 0.33 | 1 | 0.17 | 0.68 |
| $BAc$ | 1 | 0.42 | 0.52 | 1 | 0.38 | 0.54 |
| $DA \times Year$ | 2 | 28.75 | $< 10^{-5}$ | 2 | 0.35 | 0.70 |
| $DA \times Site$ | 1 | 0.78 | 0.38 | 1 | 2.56 | 0.11 |
| $PA \times Year$ | 2 | 1.35 | 0.27 | 2 | 2.01 | 0.14 |
| $PA \times Site$ | 1 | 4.11 | 0.05 | 1 | 0.29 | 0.59 |
| $BAc \times Year$ | 2 | 4.39 | 0.02 | 2 | 2.04 | 0.14 |
| $BAc \times Site$ | 1 | $< 10^{-3}$ | 0.98 | 1 | 1.67 | 0.20 |
| $Year \times Site$ | 2 | 12.61 | $< 10^{-4}$ | 2 | 7.50 | $< 10^{-2}$ |
| LMMs fitted with the AUDPC of DM and PM on the clusters | | | | | | |
| Expl. var. | NumDf | F value | Pr(>F) | NumDf | F value | Pr(>F) |
| $Year$ | 2 | 4.04 | 0.02 | 2 | 18.93 | $< 10^{-5}$ |
| $Site$ | 1 | 11.07 | $< 10^{-2}$ | 1 | 10.66 | $< 10^{-2}$ |
| $DAc$ | 1 | 23.49 | $< 10^{-4}$ | 1 | 3.79 | 0.06 |
| $PAc$ | 1 | 5.05 | 0.03 | 1 | 0.38 | 0.54 |
| $BAc$ | 1 | 0.04 | 0.84 | 1 | 0.20 | 0.66 |
| $DAc \times Year$ | 2 | 11.58 | $< 10^{-4}$ | 2 | 1.53 | 0.22 |
| $DAc \times Site$ | 1 | 6.02 | 0.02 | 1 | 0.32 | 0.57 |
| $PAc \times Year$ | 2 | 0.24 | 0.79 | 2 | 1.53 | 0.22 |
| $PAc \times Site$ | 1 | 0.99 | 0.32 | 1 | 1.16 | 0.28 |
| $BAc \times Year$ | 2 | 1.06 | 0.35 | 2 | 0.66 | 0.52 |
| $BAc \times Site$ | 1 | 0.28 | 0.60 | 1 | 0.68 | 0.41 |
| $Year \times Site$ | 2 | 10.69 | $< 10^{-3}$ | 2 | 5.33 | 0.01 |

Importantly, our results on the oenological quality of the harvest were derived from analyzing musts made exclusively from healthy berries. Accordingly, this approach reveals the indirect effects of DM on the foliage or berries on harvest quality after sorting out diseased berries. The potential alcohol content was reduced by up to 2% vol with increased DM severity (Figure S3.4 A). Similarly, Jermini et al. (2010) observed a loss of yield quality, more specifically a reduction of total soluble solids in berries, with increasing DM severity as generated by their "standard schedule" (low DM severity) and "untreated canopy" (high DM severity) modalities. In contrast to yield, the determinants of $AlDg$ appeared more complex since the effect of DM severity on both the foliage and the clusters is varying with sites and/or the years. An increase of the severity of DM on the foliage of 1% decreases $AlDg$ by 0.18 % vol in 2006, 0.06 % vol in 2007 and 0.014% vol in 2008. The greater impact of DM in 2006 may be attributed to a later onset of DM infection, primarily occurring outside the period of bunch sensitivity, compared to 2007 and 2008. Consequently, the ratio of leaf surface area to fruit mass, the amount of carbohydrates accumulated, and their conversion to sugar in the must (which mainly occurred after crop stage 81) were all lower in 2006 (Naor et al., 2002). Since higher sugar content leads to higher potential alcohol content (Ribéreau-Gayon et al., 1998), late DM infections are likely to result in reduced alcohol content. In contrast, $pH$ is not significantly affected by changes in the leaf-to-fruit ratio (Parker et al., 2015), which may explain the weak effect of DM severity on the foliage and the absence of effect on the clusters observed in our study.

The results also indicated slightly significant effect of interactions between $PA \times Site$ (p-value=0.05) and $BAc \times Year$ (p-value=0.02) for the potential alcohol $AlDg$, as well as slightly significant direct effect of $PAc$. As outlined, BOT epidemics developed very little during the experiments, even in the plots without fungicide application. The standardized AUDPC of BOT always remains lower than 0.5%, making it impossible to study the effect of BOT on qualitative yield. However, it is important to note that a low level of visible symptoms of Botrytis bunch rot can be misleading as symptomless latent infections can also impact qualitative yield (Fedele et al., 2020). Similarly, the epidemics of PM was almost weak in our experimental desgin. The mean standardized AUDPC of PM severity on clusters was always below 2% for all the factor combinations considered, except at Latresne in 2006 where it reached 4.4 % without fungicide control. In the literature, the threshold of 3% of infected berries is mentioned as the level at which off-flavors become perceptible in wine (Ough & Berg, 1979). More specifically, bunches affected by powdery mildew tend to exhibit reduced levels of total soluble solids (Gadoury et al., 2001; Stummer et al., 2003), resulting in lower potential alcohol. The low variability in PM severity within our experimental design limited our ability to go a step further and determine the threshold at which infected berries begin to affect total soluble solids and other qualitative indicators.

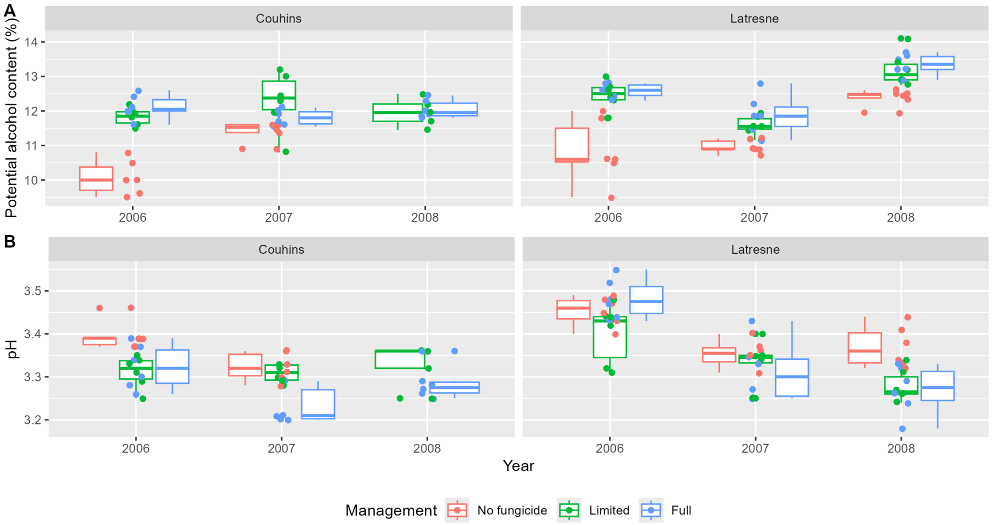


**Figure S3.1. Qualitative yield variables.** (A) Potential alcohol content measured at harvest. Each boxplot is based on the 6 measures done for each combination of site (Couhins and Latresne), cropping seasons (2006 to 2008) and fungicide managements levels (no, limited and continuous). (B) Same as (A) for the pH.

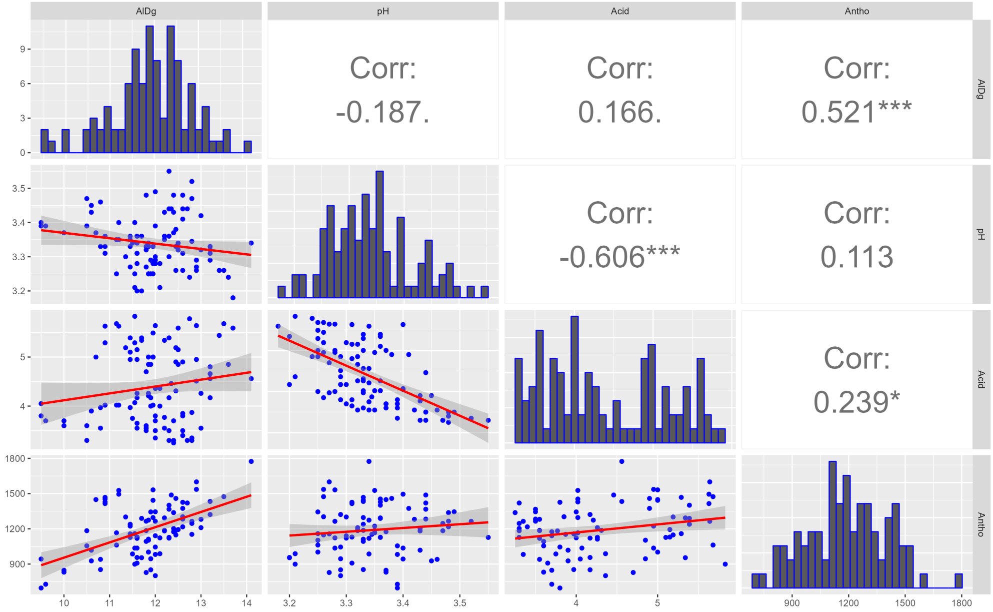


**Figure S3.2. Correlation matrix between the four quality variables**. The correlation matrix and the pairwise scatter plots were computed on 101 observations with R package *ggcorr* between the four yield quality variables ($AlDg$, $pH$, $Acid$ and $Antho$).

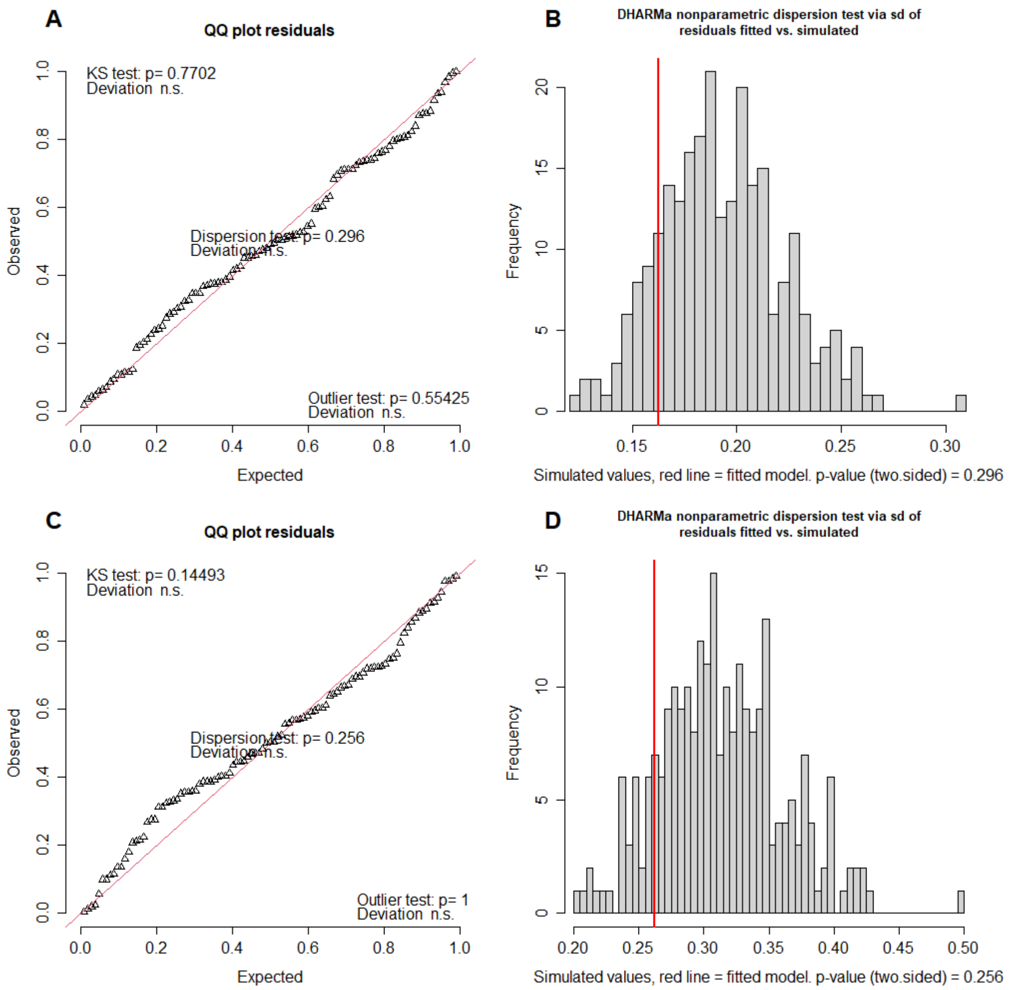


**Figure S3.3. Residual plots for the LMM fitted to the response variable potential alcohol content**. A,B: Quantile-quantile plot (A) and Simulation-based dispersion test (B) for the LMM on $AlDg$ with the AUDPC of PM and DM on the foliage (variables $PA$ and $DA$). C,D: Same as panels A and B for the LMM on $AlDg$ with the AUDPC of PM and DM on the clusters (variables $PAc$ and $DAc$). The plots are realized using the R package DHARMa.

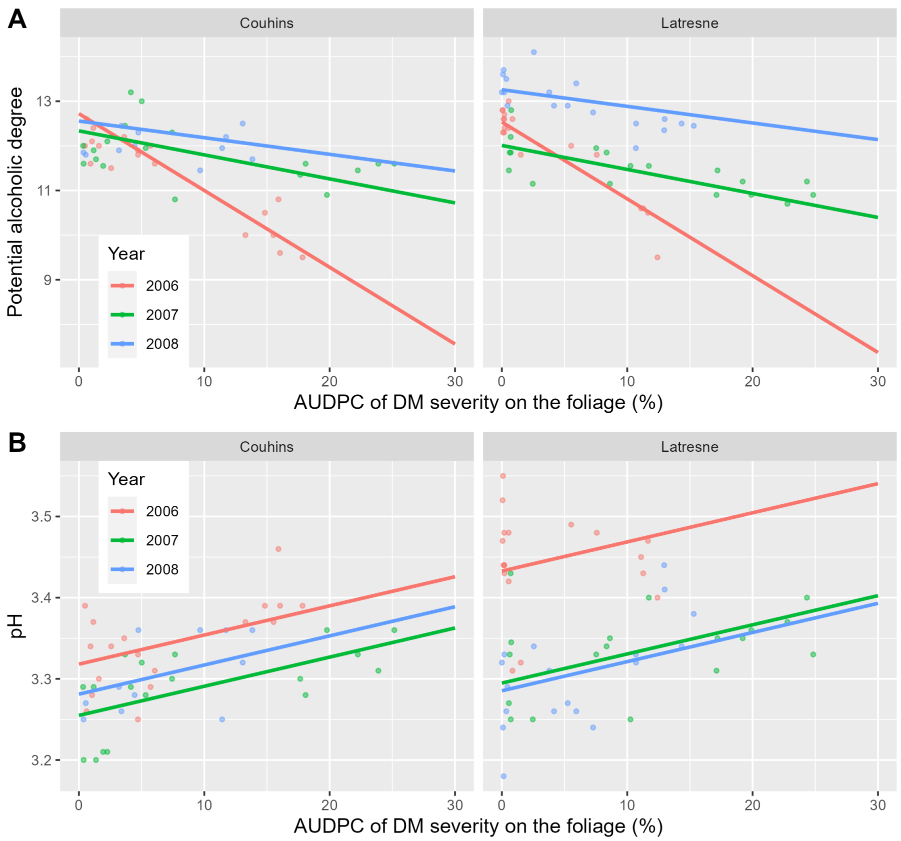


**Figure S3.4. Relationships between downy mildew severity on the foliage and yield quality variables.** (A) Potential alcohol content (response variable $AlDg$) as a function of the AUDPC of DM on the foliage fitted with the LMM. (B) Same as (A) for the response variable $pH$. The curves of model fits were realized by considering only the significant effects in the LMMs. For $AlDg$, the values of $PA$ and $BAc$ were set to their mean values 1.48 and 0.08. The full line represents the mean estimates, and the points the experimental measures.

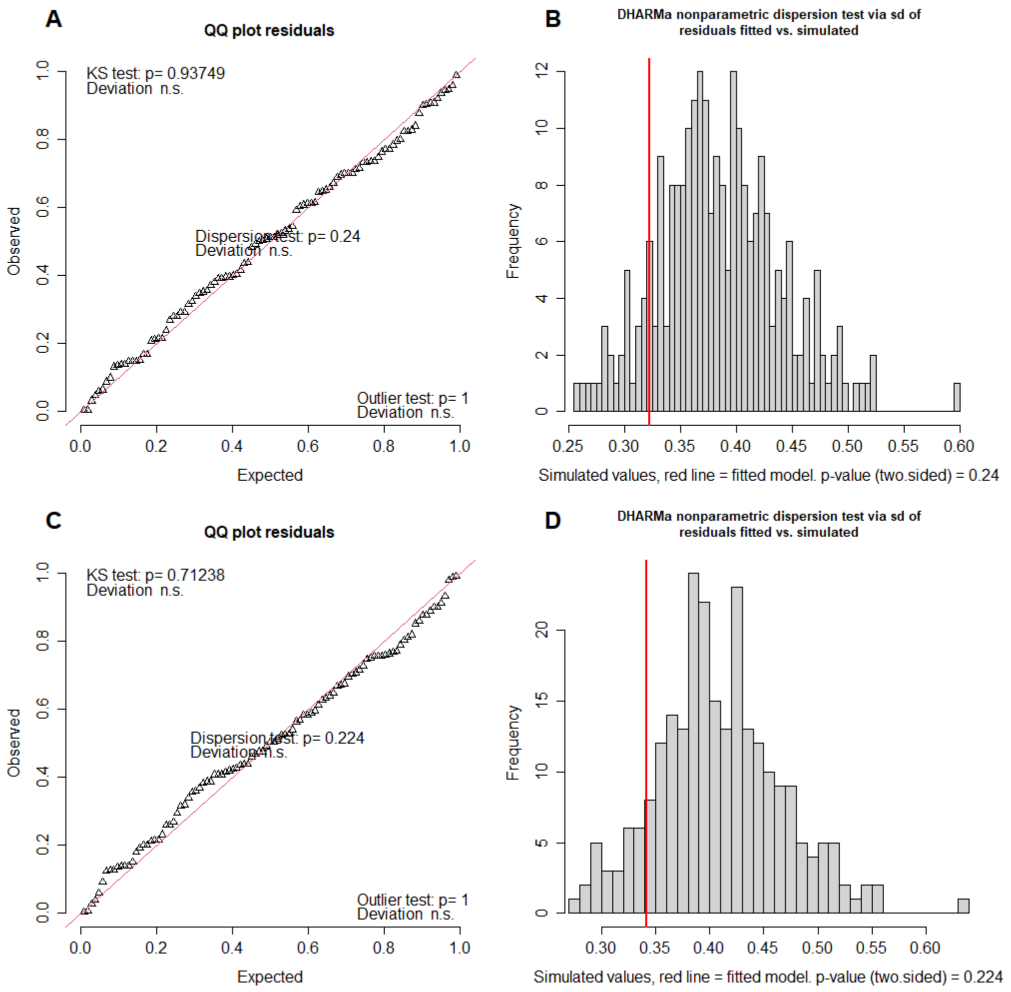


**Figure S3.5. Residual plots for the LMM fitted to the response variable pH.** (A,B) Quantile-quantile plot (A) and Simulation-based dispersion test (B) for the LMM on $pH$ with the AUDPC of PM and DM on the foliage (variables $PA$ and $DA$). (C,D) Same as panels A and B for the LMM on $pH$ with the AUDPC of PM and DM on the clusters (variables $PAc$ and $DAc$). The plots are realized using the R package DHARMa.
